# *O*-GlcNAcylation effect on Tau in modulating its seeding and cellular transmission in Alzheimer’s Disease

**DOI:** 10.64898/2026.05.27.728078

**Authors:** Debapriya Kundu, Wei-Wei Chang, Wan-Chen Lu, Lee-Way Jin, Wan-Chen Huang, Yun-Ru Chen

## Abstract

Post-translational modifications critically regulate neurodegenerative disease progression. In Alzheimer’s disease (AD) and tauopathies, Tau hyperphosphorylation promotes aggregation and pathological spreading, whereas *O-*GlcNAcylation has emerged as a protective modification alongside its interplay with phosphorylation. However, the role of in vitro site-specific and global *O*-*G*lcNAcylation in Tau proteinopathy remains elusive. Here, using full-length Tau-441 (2N4R), we showed that *O*-GlcNAcylated Tau aggregated slowly, formed distinct aggregate morphology and exhibited reduced seeding capacity compared to wild-type (WT) Tau. Under phase-separated conditions, *O*-GlcNAcylated Tau formed oligomer like condensates. Mutation of the key *O*-GlcNAc sites reduced *O*-GlcNAc-transferase (OGT) mediated *O*-GlcNAcylation, cellular transmission and affected cross-talk with phosphorylation relative to WT. OGT overexpression alleviated WT-Tau toxicity in cells, and O-Tau fibrils were less toxic to primary cortical neurons compared to WT Tau fibrils. Finally, an in-house novel site-specific S422 *O*-GlcNAc-Tau antibody revealed reduced S422 *O*-GlcNAcylation in AD brain tissues, highlighting its protective role in AD pathogenesis.

## Introduction

Alzheimer’s disease (AD) which constitutes about 70 % of all dementia cases^1^ is a progressive neurodegenerative disease associated with extensive neuronal loss, brain volume shrinkage, followed by memory deficits and cognitive decline^2^. Tau pathology is one of the hallmark features of AD besides amyloid plaque^3^. Under physiological conditions, Tau binds to axonal microtubules and promotes microtubule assembly, whereas in diseased state it detaches and forms neurofibrillary tangles (NFTs), largely driven by hyperphosphorylation. Pathological Tau also spreads in a prion-like, cell-to-cell manner across neural networks in mouse and human brain^4^. In the human brain, six Tau isoforms arise from alternative splicing of MAPT gene, differing in 3R/4R repeats and N-terminal inserts. Alzheimer’s disease contains both 3R and 4R isoforms, in Pick’s disease mainly 3R, while progressive supranuclear palsy (PSP) and corticobasal degeneration (CBD) are predominantly 4R tauopathies^5^. The role of Tau in cell-to-cell transmission has been studied, where cell culture and in vivo models showed preformed synthetic Tau fibrils were able to drive seeded-propagation^6,7^. A recent study highlighted that full-length 2N4R Tau can form seed-competent intraneuronal inclusions in humanized mouse cortical neuronal model^8^. Tau undergoes multiple post translational modifications (PTMs), which regulate its structure, localization, interactions, and aggregation^9^, with phosphorylation being most implicated and extensively studied in AD and other tauopathies^10^. Another Tau PTM a Serine/Threonine dynamic modification of O-linked-β-N-acetyl-D-glucosamine (*O*-GlcNAc)^11^, has been shown to inhibit α-synuclein aggregation, toxic oligomer and fibril formation, and in vivo seeding, thereby suppressing Parkinson’s disease pathogenesis^12–14^.

*O*-GlcNAc is an O-linked glycosylation which is the addition of a single molecule of monosaccharide N-acetylglucosamine on the hydroxyl (-OH) group of the Ser/Thr residues^11,15^. Unlike protein phosphorylation and dephosphorylation that are mediated by multiple kinases and phosphatases respectively, *O*-GlcNAcylation acted as an intracellular sensor for glucose assimilation^16^, is regulated by only two enzymes, namely *O*-GlcNAc transferase (OGT) and *O*-GlcNAcase (OGA). OGT catalytically adds *O*-GlcNAc moiety to naked Ser/Thr residues from its precursor uridine diphosphate N-acetylglucosamine (UDP-GlcNAc), while, the reversible removal was done by OGA^17^. *O*-GlcNAcylation has several implications in AD. Previous literature showed *O*-GlcNAcylation regulated Tau phosphorylation and affected the extent of neurodegeneration^11,18^. Knockdown of OGT downregulated *O-*GlcNAcylation of Tau, which in turn upregulated phosphorylation, modulated by glucose metabolism in HEK293 cells, rodent and AD brain^19^. The treatment of OGA inhibitors increased global *O*-GlcNAcylation and decreased pathological Tau in various transgenic AD mice models^20–22^. Also, *O*-GlcNAcylation at a prominent site (Ser400), on Tau directly inhibited its oligomerization, without altering its global structure^23^.

Although *O*-GlcNAcylation of Tau was studied in vivo, there lies a vacuum regarding its role in in vitro seeding and cellular transmission. Hence, in this study we aim to investigate and elucidate the role of global and site-specific Tau *O*-GlcNAcylation for aggregation, seeding, cellular transmission, cross-talk and toxicity by using full-length 2N4R (Tau-441) with or without *O*-GlcNAcylation. We first expressed and purified recombinant wild type (WT) Tau-441 and its *O*-GlcNAcylated form (O-Tau) in *E. coli* and examined their aggregation, seeding, and diffusion dynamics under phase separated (LLPS) condition. We identified critical *O*-GlcNAc sites on Tau and studied their effect on cellular transmission, cross-talk relationship with phosphorylation, cytotoxicity and neuronal toxicity. Our results collectively demonstrated that *O*-GlcNAcylation altered the aggregation propensity by forming soluble, small oligomeric assemblies, which are less toxic and increase cellular transmission. Furthermore, we generated an in-house *O*-GlcNAc-Tau antibody at the S422 site which detected *O*-GlcNAcylation both in vitro and in vivo post mortem AD patient tissues.

## Results

### Characterization and determination of *O*-GlcNAcylation on full-length Tau-441

To characterize WT and *O*-GlcNAcylated Tau-441, recombinant WT and *O*-GlcNAc-Tau-441 (O-Tau) were expressed and purified from *E. coli* BL21 cells. WT was expressed and purified following previous literature^24^. O-Tau was generated under co-expression of WT and OGT, purified firstly by SP Sepharose cation-exchange column and secondly by reverse phase (RP)-HPLC, followed by *O*-GlcNAcylation detection. as shown schematically in Figure S1A. The RP-HPLC chromatogram revealed three elution peaks P1, P2, P3 of O-Tau, with P1 showing highest intensity (Figure S1B). The elution fractions were analyzed firstly by dot-blot analysis using an anti-S400-*O*-GlcNAc-Tau specific antibody (O-Tau-S400) detecting Tau *O-*GlcNAcylation at S400 site^25^ and by RL2, a pan specific *O*-GlcNAc antibody^11^ (Figure S1C). *O*-GlcNAcylation was detected in fractions P1, P2 and the O-Tau SP sepharose fraction with both antibodies, whereas purified WT-Tau showed no detectable signal, confirming no *O*-GlcNAcylation. Both WT and O-Tau fractions were positive for total Tau antibody Tau-5. These elution fractions were further analyzed by LC-MS to detect total protein molecular masses (Figure S2A and S2B). Deconvolution peak analysis showed P1 was a predominantly monomeric peak having 100% total intensity, with average molecular mass of 46771.7 Da for ∼4-5 *O*-GlcNAc moieties per Tau molecule (Figure S2A). P2 was predominantly a dimeric peak of average mass 46779 Da (43.75) % total and 93558.9 Da (24.37) % total intensities respectively (Figure S2B). MALDI-TOF analysis further confirmed P1 having a monomeric peak of high relative intensity at 46363 Da, (Figure S3A), while P2 having a dimeric peak at 93351.2 Da, and P3 primarily noise (Figures S3B and S3C). This further reiterates that the peaks obtained from RP-HPLC had different *O*-GlcNAc populations. While P1 had a relatively monomeric population of *O*-GlcNAcylated Tau protein, P2 had a dimeric population.

Additionally, to confirm *O*-GlcNAc sites, the monomeric P1 fraction was further trypsin digested and analyzed by LC-MS/MS technique. Several *O*-GlcNAc sites, with number of identifications i.e., T111, S131, S400, S409, N410, S412, S413 S416, S422, were shown in (Figure1A). Among these sites S400, S412, S413, S422 matched with previously reported *O*-GlcNAc sites^26,27^. Most identified sites were located within the C-terminal domain of Tau. In contrast, WT Tau showed a single RP-HPLC elution peak under identical conditions (Figure S4A) and a molecular mass of ∼45.5 kDa by MALDI-TOF analysis (Figure S4B), confirming the absence of modification. Together, these results demonstrate the successful generation and purification of predominantly monomeric *O*-GlcNAcylated Tau-441 suitable for downstream functional studies.

### *O*-GlcNAcylation of Tau reduced its aggregation propensity

The fibrillization profile of WT and *O*-GlcNAc–Tau (O-Tau) was studied in the presence of heparin using Thioflavin T(ThT) fibrillization assay, which monitors amyloid cross-β-sheet formation through enhanced fluorescence^28^. We first centrifuged purified WT Tau and *O*-Tau at 17,000 g to remove preformed aggregates, followed by the ThT fluorescence analysis. The results showed that WT fibrillized faster than its *O*-GlcNAcylated form with higher final ThT intensity, whereas O-Tau aggregated with a slower elongation phase (Figure 1B). The end-point products of ThT assay were subjected to transmission electron microscopy (TEM) imaging to observe the morphology and Fourier-transform infrared (FTIR) spectroscopy to examine the secondary structure. The TEM images (Figure 1C) showed that WT Tau formed longer and well-extended fibrils averaging at ∼1191 nm whereas O-Tau formed much shorter fibrils averaging at ∼206 nm, and some amorphous aggregates. The average fibril lengths were determined from nine fibrils across three TEM images. The FTIR spectra (Figure 1D) showed strong and prominent β-sheet peak at 1633 cm^-1^ for WT^29^, and a very minor shoulder at 1657 cm^-1^. O-Tau had a broader peak, with maxima at 1641 cm^-1^ indicating disordered or random-coil like morphology, with a minor β-turn like shoulder at 1675 cm^-1^. The FTIR data were summarized in Table 1. Together, our results showed that *O*-GlcNAcylation reduces Tau fibrillization propensity and alters the structure and fibrillar morphology.

**Figure 1.**
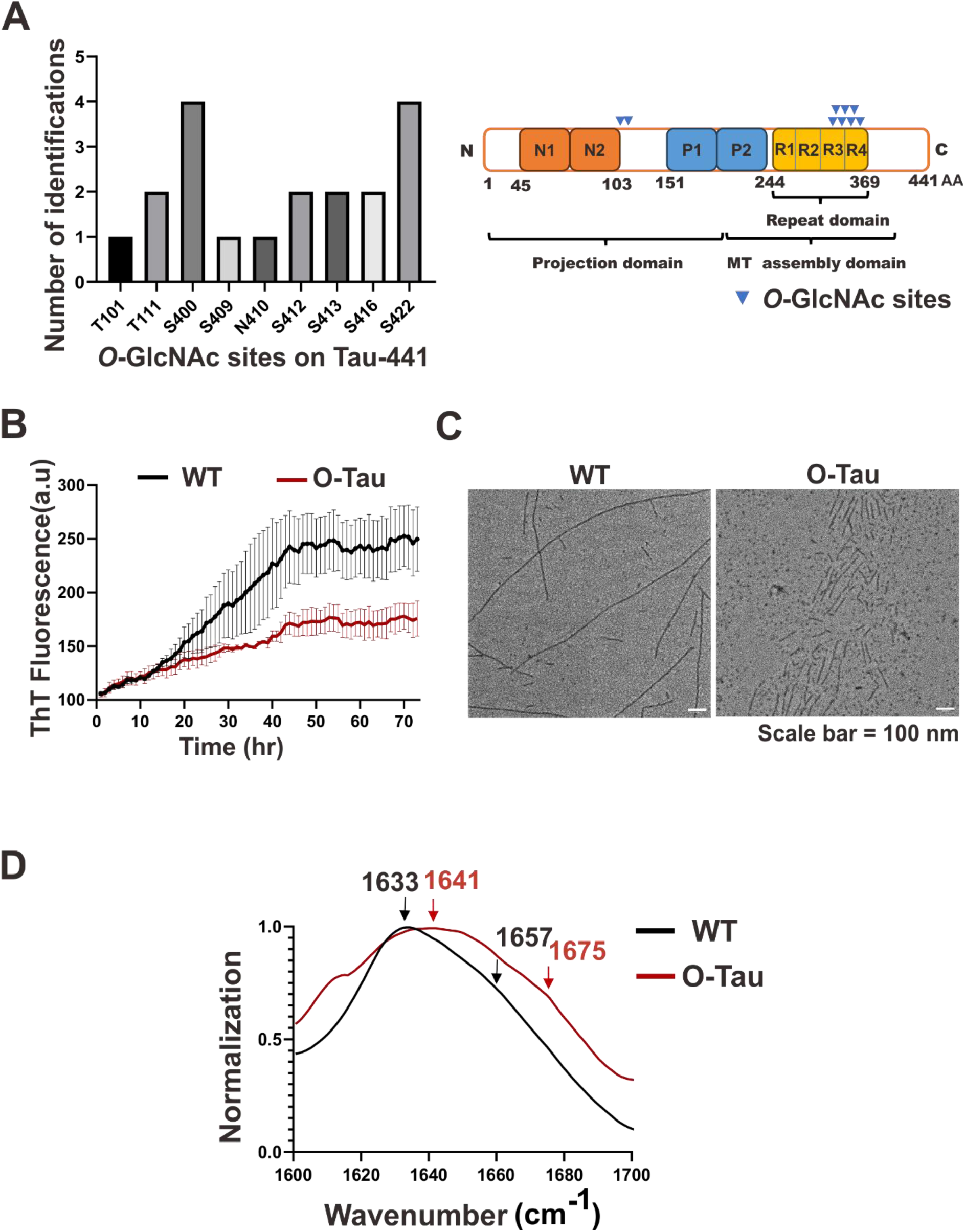
*O*-GlcNAc site determination on FL-Tau-441 and ThT fibrillization of WT and O-Tau. **(A)** Graphical and pictorial representation of *O*-GlcNAc sites identification by LC-MS/MS analysis from four independent repeats (N=4). Several *O*-GlcNAc sites, i.e., T111, S131, S400, S409, N410, S412, S413 S416, S422, were detected. **(B)** ThT fibrillization assay of WT and O-Tau monomers, induced by heparin, to form fibrils. **(C)** TEM images of WT and O-Tau fibrils after ThT fibrillization, respectively (scale bar = 100 nm). **(D)** FTIR spectra of WT and O-Tau fibrils after ThT fibrillization. FTIR data between 1600 and 1700 cm^-1^ were collected and normalized. All experiments were performed in triplicates.

**Table 1.** FTIR peak positions of Tau fibrils.

| **Tau Samples** | **α-helix (cm<sup>-1</sup>)** | **β-sheet (cm<sup>-1</sup>)** | **β-turn (cm<sup>-1</sup>)** | **Random coil /**<br>**unordered (cm<sup>-1</sup>)** |
| --- | --- | --- | --- | --- |
| WT + Heparin | 1657 | 1633 |  |  |
| O-Tau + Heparin |  |  | 1675 | 1641 |
| WT + WT seed | 1659 | 1631 |  |  |
| WT + O-Tau seed |  |  |  | 1637, 1650 |
| O-Tau + WT seed |  | 1632 | 1667 |  |
| O-Tau + O-Tau seed |  |  | 1667 | 1638 |

### *O*-GlcNAc-Tau exhibited lower seeding capability

Next, we examined the seeding properties of O-Tau. Seeding is a general property for amyloids. By addition of preformed fibril seeds as a nucleus, the fibrillization escapes the nucleation process and directly goes into elongation phase^30^. The fibril seeds were prepared by sonicating the end-point products of ThT assay. WT and O-Tau monomers were incubated with or without 5% WT or O-Tau fibril seeds in the presence of heparin in the ThT assay (Figure 2A). The WT seeds exhibited higher seeding effect compared to O-Tau seeds. Seeding effect in WT monomer with WT seeds was characterized by an intense and thick fibrillization and sharp decrease in lag phase time from 11 hr to ∼ 3 hr. In contrast, O-Tau seeds could not seed the WT monomer as effectively as WT seeds. The result showed homotypic seeding in WT is stronger than heterotypic seeding. Next, O-Tau monomers were incubated with or without 5% WT or O-Tau fibril seeds under the same conditions. Interestingly, WT seeds promoted stronger seeding of O-Tau monomer than O-Tau seeds. The lag phase of O-Tau alone (∼11 hr), decreased to ∼3 hr in the presence of WT seeds, followed by an increase in elongation phase. Addition of O-Tau seeds to O-Tau monomer also reduced the lag phase, but to a lesser extent compared to the WT seeds (∼8 hr). The result indicated that O-Tau seeds were less efficient than WT seeds in seeding O-Tau monomer (Figure 2B).

**Figure 2.**
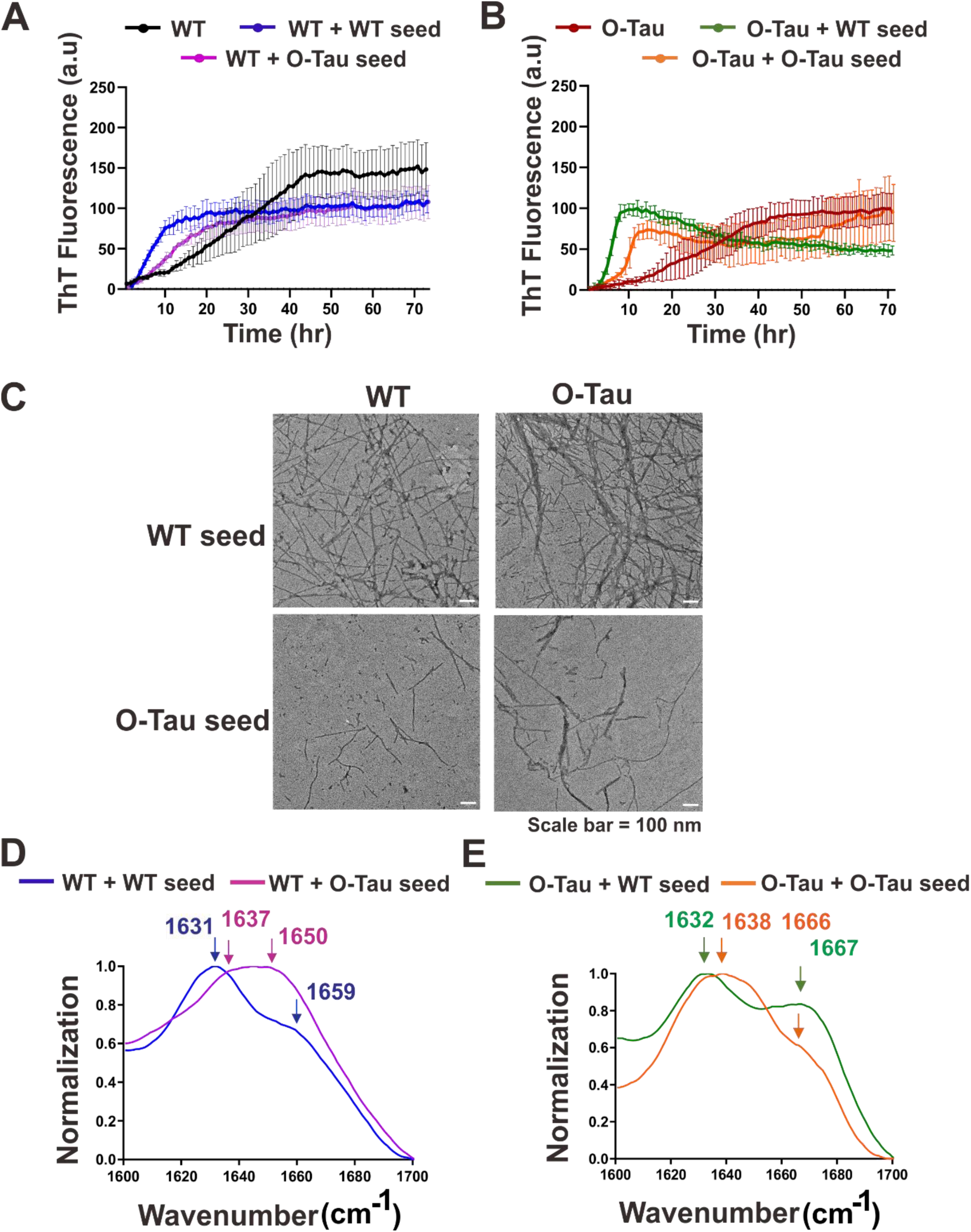
ThT seeding assay for WT and O-Tau induced by Tau preformed fibrils. **(A-B) WT (A)** and O-Tau **(B)** monomers were incubated with or without WT or O-Tau preformed fibrils. ThT fluorescence intensity was determined in the presence of heparin. Signals were obtained after subtraction from the buffer background. **(C)** TEM imaging of the seeded fibrils. Morphology of the end-point products of ThT seeding assay were collected and imaged by TEM (scale bar, 100 nm). **(D-E)** FTIR spectroscopic analysis for WT ± seeds **(D)** and O-Tau ± seeds **(E)**. Spectra was collected from 1600 and 1700 cm^-1^ and normalized. Experiments were performed in triplicates.

The morphology and secondary structure of the end-point products were further characterized using TEM and FTIR, respectively. TEM images, showed that both WT and O-Tau monomer formed abundant long fibrils when seeded with the WT seeds (Figure 2C), but O-Tau seeds addition resulted in shorter fibril structure with lower fibrillar density. In the FTIR spectra, WT monomer in the presence of WT seeds showed a sharp peak at 1631 cm^-1^ for ß-sheet structure and a minor shoulder at 1659 cm^-1^ with some α-helical characteristics, but in the presence of O-Tau seeds, the spectra predominantly shifted towards more disordered-like species having a broader peak with maxima at 1650 cm^-1^ and a minor shoulder at 1637 cm^-1^ (Figure 2D). On the other hand, WT seeds addition to O-Tau monomer formed more WT-like spectra having a mixture of β-sheet at 1632 cm^-1^ and a prominent shoulder at 1667 cm^-1^ for β-turn. In contrast, O-Tau seeds with O-Tau monomer generated a predominantly O-Tau like spectra, having a broad peak of random coil like morphology at 1638 cm^-1^, and a minor shoulder at 1667 cm^-1^ for β-turn (Figure 2E). Together, these results demonstrated that WT seeding resulted in more structured WT-like fibrils both in WT and O-Tau monomers, whereas O-Tau fibrils displayed reduced seeding effect. The FTIR spectra is summarized in Table 1.

### *O*-GlcNAcylation of Tau altered its diffusion dynamics

Tau is an intrinsically disordered protein that can undergo complex coacervation or LLPS under crowded conditions^31^. The electrostatic charge distribution in Tau, along with the aggregation prone repeat domains, can trigger liquid droplet formation under crowded condition^32–34^, as diagrammatically show in Figure 3A. Hence, we examined the LLPS properties of WT Tau and O-Tau. WT and O-Tau monomers were fluorescently labelled with Alexa flour 488 dye and the in vitro condensate formation was examined by confocal fluorescence microscopy in the presence of a crowding agent, PEG 8000. Previously, it has been shown that Tau undergoes LLPS under 10% PEG 8000 in NaCl buffer condition. Hence, droplet formation of 10 µM WT and O-Tau was examined with increasing NaCl concentrations under 10% PEG 8000 (Figure S5A). Comparison of droplet area and diameter (Figure S5B and S5C), showed no significant difference between WT and O-Tau, although *O*-GlcNAcylation showed a trend toward larger droplets. Next, we used fluorescence recovery after photobleaching (FRAP) technique to investigate protein dynamics within the droplets. WT and O-Tau were subjected to FRAP at 10 min and 5 hr post droplet formation. At 10 min both WT and O-Tau showed ∼70% fluorescence recovery, with no significant difference in final intensity (Figure 3B). Interestingly, when FRAP was conducted at 5 hr after induction, neither WT nor O-Tau demonstrated observable recovery (Figure 3C), exhibiting transition toward a gel-like state. We further examined the diffusion coefficients of WT and O-Tau under LLPS conditions using fluorescence correlation spectroscopy (FCS).

**Figure 3.**
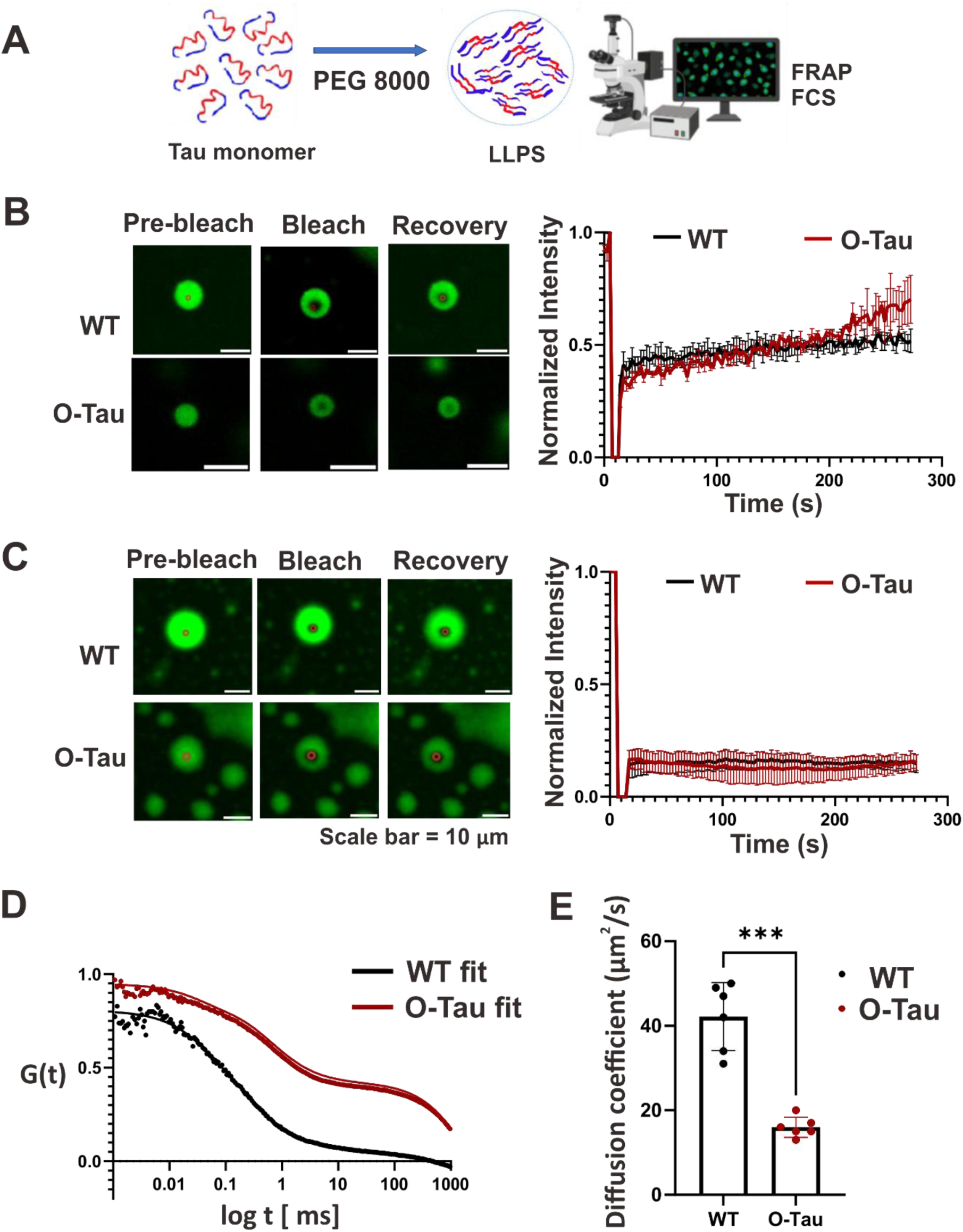
Determination of Tau droplet dynamics by FRAP and FCS. **(A)** Schematic representation of Tau LLPS induced by the crowding reagent, PEG 8000, followed by FRAP and FCS analyses. **(B-C)** FRAP analysis of WT and O-Tau droplets at 10 min **(B)** and 5 hr **(C)** after droplet induction. Droplets were formed in HEPES buffer, pH7.4, containing 50 mM NaCl and 10% PEG 8000. Fluorescence recovery was monitored for 270 s after photobleaching in three independent replicates. **(D-E)** FCS analysis of Tau diffusion within condensates. Autocorrelation curves are shown in **(D)**, and calculated diffusion coefficients are shown in **(E).** Condensates were formed using 10 µM unlabeled WT or O-Tau monomers doped with 50 nM Alexa Flour-488 labeled WT or O-Tau. Data are presented as mean ± (s.d.); n = 6 droplets per condition. Statistical significance was determined by Student’s t-test. \*\*\**P* < 0.001.

Hydrodynamic radii were estimated using the Stokes-Einstein equation, allowing approximate estimation of their molecular masses. Diffusion parameters were obtained by fitting auto-correlation curves to a two-component diffusion model with triplet (3D) state. We plotted the average curve fits of auto-correlation functions vs diffusion time, for 6 WT (black, n=6) and O-Tau (red, n=6) condensates, (Figure 3D). WT Tau diffused faster and was well described by a single homogenous species, whereas O-Tau diffused more slowly and exhibited two different diffusing species, indicating the presence of heterogeneous assemblies. The average diffusion coefficient (D) of WT Tau corresponded to 42.2 ± 6.933 µm²/s, while O-Tau displayed reduced diffusion coefficient of 16.0 ± 0.925 µm²/s (Figure 3E). Based on the Stokes–Einstein equation, R_h_ = K_b_T/6πηD^35^, where the hydrodynamic radius (R_h_) and the diffusion coefficient are inversely related, the relative R_h_ of O-Tau was ∼2.6-fold larger than WT. Using polymer scaling laws for intrinsically disordered proteins (IDPs), where molecular mass is correlated with R_h_ as M ∝ *Rh*^1⁄v^ (with, v ∼ 0.57 for IDPs^36^), the apparent molecular mass of O-Tau was estimated to be ∼5.5-fold larger than WT. Assuming the WT species corresponds to the expected Tau-441 monomer mass (∼45.5 kDa), the apparent molecular mass of O-Tau was calculated at ∼249.1 kDa, consistent with a small oligomeric assembly comprising ∼5–6 Tau subunits.

To further investigate whether *O*-GlcNAcylation modulates Tau protein interactions in live HEK293 cells, we performed fluorescence lifetime imaging microscopy-FRET (FILM-FRET) analysis. EGFP-Tau and mCherry-Tau (either WT or the S400A/S422A double mutant) were co-expressed with or without OGT overexpression (Figure S6A). FRET efficiency was quantified based on donor fluorescence lifetime measurements (Figure S6B). The double mutant exhibited a significantly higher FRET efficiency than the WT, indicating altered intermolecular arrangement or enhanced self-association. Co-expression with OGT significantly increased the FRET efficiency of WT Tau but no effect on the double mutant (Figure S6C). These findings suggest that *O*-GlcNAcylation promotes closer intermolecular association and structural proximity of Tau.

### OGT overexpression resulted in higher *O*-GlcNAcylation in WT but not in *O*-GlcNAc-Tau mutants

We focused on the two critical *O*-GlcNAc sites, namely S400 and the S422, that showed higher frequency of appearance during LC-MS/MS analysis, for further examination. We validated the importance of these *O*-GlcNAc sites of Tau in HEK293 cells. Both EGFP- and mCherry fused Tau (gT or mT, respectively) with single and double mutants for the two *O*-GlcNAc sites (S400 and S422) to alanine were prepared by site-directed mutagenesis. The single and double mutants were transiently transfected with OGT overexpression in HEK293 cells. The cell lysate was further collected, followed by immunoprecipitation with the total Tau antibody Tau-5. *O*-GlcNAc and total Tau levels were detected by Western blot analysis (Figure 4A) and quantified (Figure 4B), respectively. The results showed that the normalized *O*-GlcNAcylated Tau level in the two single mutants, ie, gT-S400A and gT-S422A, was substantially decreased. For the double mutant gT-S400A/S422A, the decrease in *O*-GlcNAc level was similar to gT-S442A. Previous literature has shown S400 as an important *O*-GlcNAc site in vitro^37^, our data further reiterates its importance. The chorological decrease in *O*-GlcNAc signals for the gT-S422A mutant, and for the double mutant gT-S400A/S422A further sheds light on the importance of S422 also as a prominent *O*-GlcNAc site, apart from the S400 site.

**Figure 4.**
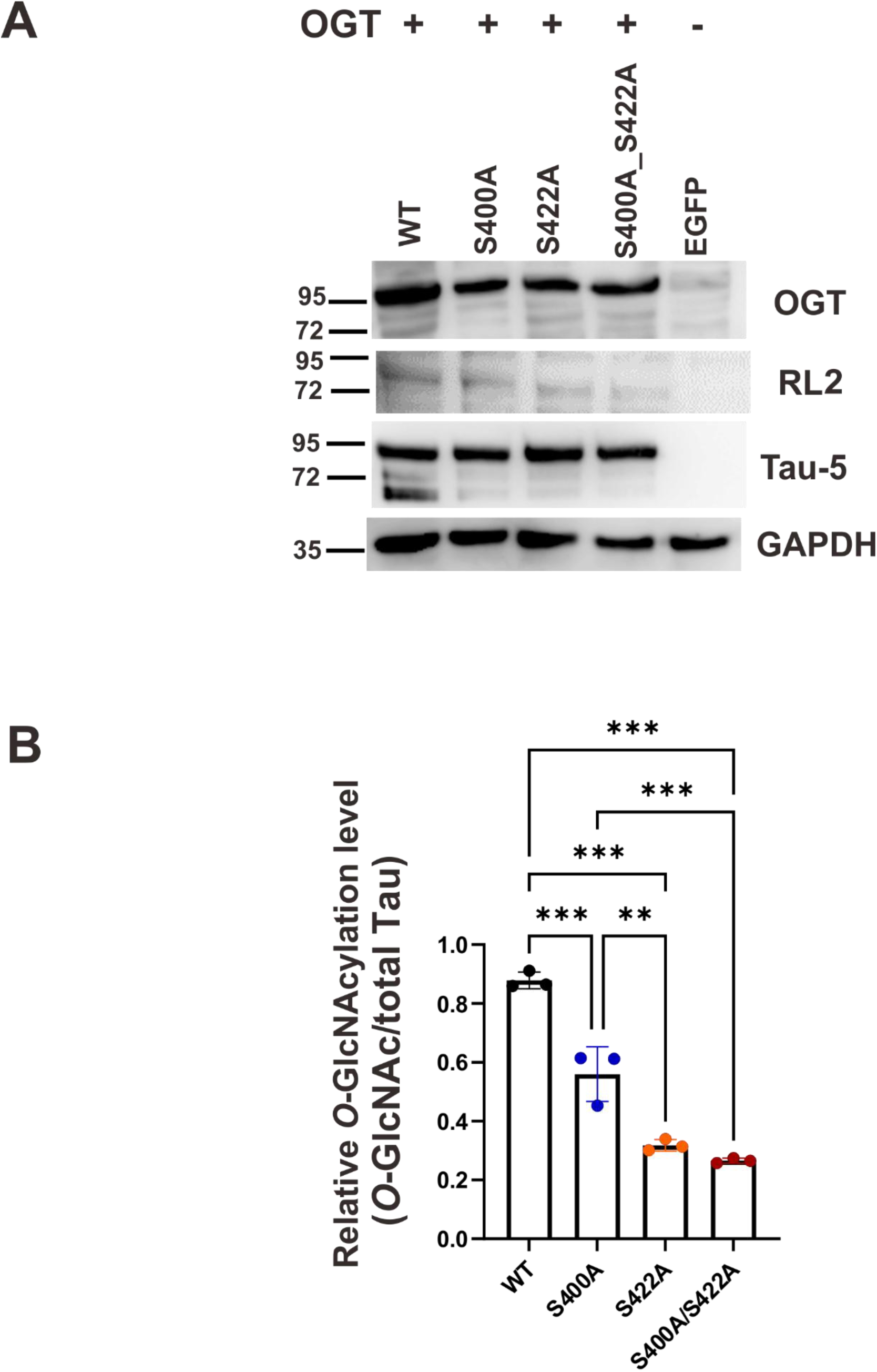
Determination of *O*-GlcNAcylation expression level of WT and *O*-GlcNAc-Tau variants in cells. **(A-B)** Western blot analysis **(A)** and quantification **(B)** of EGFP-Tau-441 (gT-WT), EGFP-Tau-441-S400A (gT-S400A), EGFP-Tau-441-S422A (gT-S422A), EGFP-Tau-441-S400A/S422A (gT-S400A/S422A), in the presence of OGT overexpression. The proteins were overexpressed in HEK293 cells for 3 days followed by immunoprecipitation and western blot detection of *O*-GlcNAc and total Tau levels using RL2 and Tau-5 antibodies, respectively. EGFP alone was used as a negative control. Relative *O*-GlcNAcylation levels were quantified as the ratio of *O*-GlcNAc to total Tau level from 3 independent replicates using ImageJ analysis. Statistical significance was calculated by one-way ANOVA, with Tukey’s multiple comparison post hoc test. \**P* < 0.05, \*\**P* < 0.01, \*\*\**P* < 0.001.

### Tau exhibited a cross-talk relationship between *O-*GlcNAcylation and phosphorylation

Previous studies have shown that there exists a cross-talk relationship between *O*-GlcNAcylation and phosphorylation in Tau because both modifications affect similar Ser/Thr residues^38,39^. GSK-3β is one of the major kinases involved in moderate phosphorylation and hyperphosphorylation of Tau^40,41^. In order to address the cross-talk relationship, we first incubated recombinantly expressed WT and O-Tau with GSK-3β kinase in vitro and used dot-blot analysis at two phosphorylation sites S396 and S404 at the vicinity of S400 to detect if *O*-GlcNAcylation affects site-specific phosphorylation at its vicinity. Sites such as S396 and S404 were reported previously as prominent pathological phosphorylation sites in early-stage AD^42^. From dot-blot analysis (Figure 5A), WT Tau treated with GSK-3β showed both S404 and a strong S396 phosphorylation signal, as previously reported^43^. Interestingly, when O-Tau was incubated with GSK-3β no phosphorylation signal was detected at S396 and very weak phosphorylation signal at S404. The results demonstrated that *O*-GlcNAcylation on Tau at S400, reduced the phosphorylation of the proximity phosphorylation sites, when treated with GSK-3β, as previously addressed^11^. Hence, global *O*-GlcNAcylation in general and specifically at a prominent site S400 may affect site-specific phosphorylation at S396 and S404, at its vicinity which re-establishes the direct cross-talk relationship between these two post-translational modifications in recombinantly expressed Tau.

**Figure 5.**
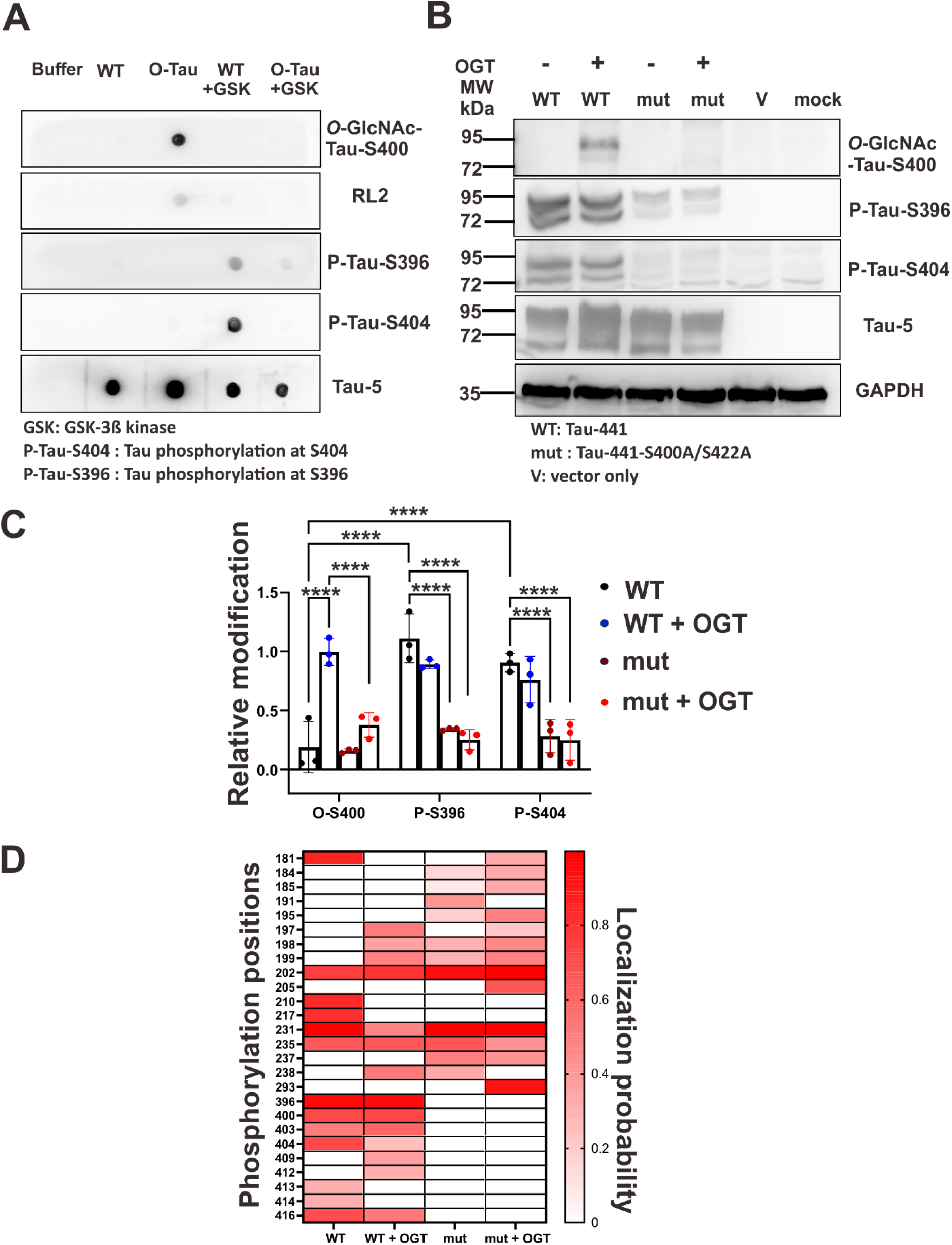
Identification of the crosstalk between *O-*GlcNAcylation and phosphorylation of Tau *in vitro*. **(A)** Dot blot analysis of recombinantly expressed WT and O-Tau with or without GSK-3ß mediated phosphorylation. The blot was probed with antibodies against site-specific Tau *O*-GlcNAcylation at S400 (*O*-GlcNAc-Tau-S400), pan-*O*-GlcNAc (RL2), site-specific Tau phosphorylation at S396 and S404, and total Tau (Tau-5). “Buffer” indicates Tris buffer used as a negative control. (**B)** Western blot analysis of mCherry-Tau-441-WT (WT) and the double *O*-GlcNAc-site mutant of Tau-441, Tau-441-S400A/S422A (mut). The Tau variants were transiently transfected into HEK293 cells with or without OGT overexpression and the cell lysate were subjected to western blotting using antibodies. The mCherry vector and cells only were used as negative controls, indicated as V and mock, respectively. **(C)** Quantification of site-specific *O*-GlcNAcylation and phosphorylation levels in Tau variants. *O*-GlcNAcylation and phosphorylation signals were quantified by ImageJ software and normalized to total Tau level. Statistical significance was determined by one-way ANOVA with Tukey’s multiple comparison post hoc test. \**P* < 0.05, \*\**P* < 0.01, \*\*\**P* < 0.001. **(D)** Heat-map representing average localization probabilities of the phosphorylation sites for (WT) and the double *O*-GlcNAc Tau mutant S400A/S422A (mut) with or without OGT overexpression in HEK293 cells, for 3 independent repeats after LC-MS/MS analysis (n=3).

To address the cross-talk relationship in cells, we examined if OGT overexpression can site-specifically impact the phosphorylation of Tau in HEK293 cells. Hence, WT-Tau and S400A/S422A double mutant were transiently transfected with or without OGT overexpression. The level of Tau *O-*GlcNAcylation at S400 and of phosphorylated Tau was quantified against total Tau at these prominent phosphorylation sites S396 and S404 at the vicinity of S400 site by Western blot (Figure 5B, C). We observe significantly higher *O*-GlcNAcylation in WT at S400 site with OGT overexpression and WT alone showed high phosphorylation at sites S396 and S404. Interestingly, the S400A/S422A double mutant showed a significant decrease in both *O*-GlcNAcylation at S400 and phosphorylation levels at sites S396 and S404 respectively, compared to the WT Tau alone with or without OGT overexpression. Total Tau levels remained unchanged in all cases. To further quantify the level of changes, we examined the localization probability values of the phosphorylation sites obtained from LC-MS/MS analysis for the WT and the S400A/S422A double mutant (Figure 5D). We observed high localization for phosphorylation events at sites spanning throughout the proline rich domain, i.e., residues 181 to 238, and the C-terminal domain, i.e., residues 396 to 416 in case of WT. With OGT overexpression we observed reduced average localization values at S396, and even more reduction in localization at S404, which are part of the C-terminal domain. Interestingly, in the case of the double mutant with or without OGT overexpression, there was a marked reduction in confidently localized phosphorylation events at the C-terminal domain. Together, these findings indicate that *O*-GlcNAcylation and phosphorylation exhibit context-dependent cross-talk, with direct inhibitory effects observed in recombinant Tau and more complex, indirect regulation in the cellular environment.

### OGT overexpression increased cell-to-cell transmission in WT-Tau compared to its *O*-GlcNAc-Tau mutants

Tau pathology and *O*-GlcNAcylation have a complex relationship. Our data has shown *O*-GlcNAcylation influenced Tau phosphorylation site-specifically at its vicinity, a critical step in Tau dysregulation and aggregation. Tau spreading through cell-to-cell propagation in the diseased brain is a key pathogenic event in neurodegenerative diseases^44^. The influence of *O*-GlcNAcylation on Tau secretion and subsequent uptake by neighboring cells leading to transmission of Tau between cells, a process linked to the spread of Tau in vitro remain elusive.

Hence, we wanted to elucidate if *O*-GlcNAcylation affect the transmission of Tau or whether alterations in OGT levels could influence Tau secretion or uptake by neighboring cells. We first overexpressed EGFP- and mCherry-Tau-WT as well as the single and double *O*-GlcNAcylation site mutants with or without OGT overexpression, and Thiamet G (TMG) administration (Figure 6). TMG, is an OGA inhibitor, that increases *O*-GlcNAcylation, has been shown to decrease AD pathology in a tauopathy mouse model^45^, and alleviate cognitive decline in Tau/APP mutant mice^46^. Here, we co-cultured equal volume of EGFP- and mCherry Tau expressing cells, followed by quantification of dual positive cells by flow cytometry as illustrated in Figure 6A. EGFP and mCherry co-culture alone was used as control. The flow cytometry plots and the quantification were shown in Figure 6B and 6C, respectively. We observed in the presence of OGT overexpression and TMG administration, there was significantly higher percentage of cells with dual fluorescence in WT Tau expressing cells. Comparing to the single and double mutants, the dual positive cells decreased, and a chronological decrease in transmission was shown from single to double mutants. The results demonstrated that *O*-GlcNAcylation significantly increased cell-to-cell transmission of WT, whereas Tau transmission was reduced in the *O*-GlcNAc Tau mutants.

**Figure 6.**
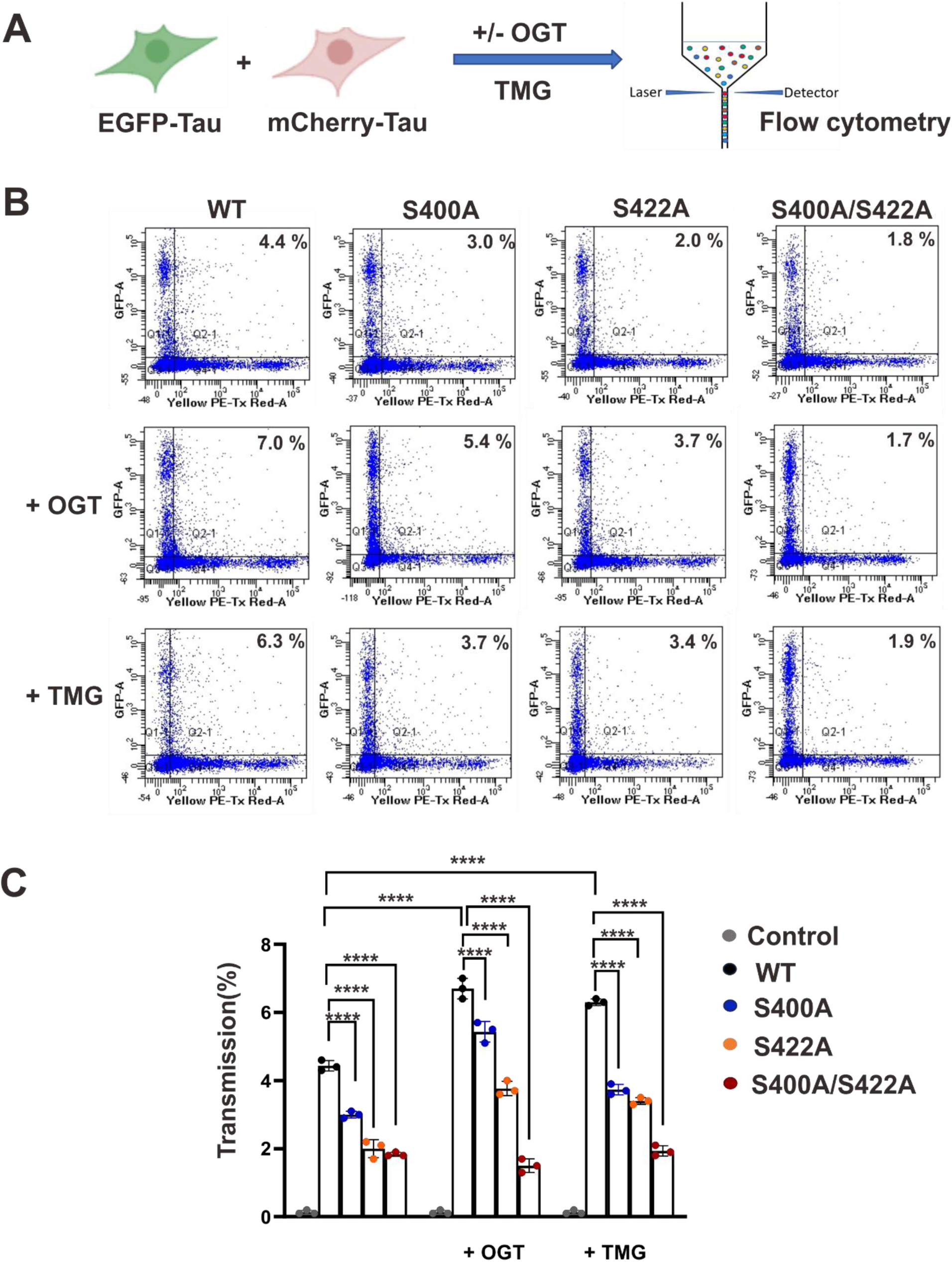
Cell-to-cell transmission analysis of WT-Tau and *O*-GlcNAc-Tau mutants. **(A)** Schematic representation of cellular transmission platform. Cells were transiently co-transfected with EGFP-Tau WT and mCherry Tau-WT single, or double mutant constructs, with or without OGT overexpression and TMG treatment. Equal number of cells were co-cultured for 48 hr, followed by flow-cytometric analysis. **(B)** Representative flow-cytometric dot plots showing EGFP and mCherry fluorescence. The percentage of EGFP/mCherry double-positive cells is indicated in each plot. (**C)** Quantification of double-positive cells as a percentage of total fluorescent cells. A total of 10,000 events were analyzed per sample for WT-Tau and *O*-GlcNAc-site Tau variants, with three replicates. Statistical significance was determined using one-way ANOVA, with Tukey’s multiple comparison post hoc test. \**P* < 0.05, \*\**P* < 0.01, \*\*\**P* < 0.001.

### Tau *O*-GlcNAcylation lowers cytotoxicity in cells and primary cortical neurons

To understand the cytotoxicity effect of Tau *O*-GlcNAcylation in cells, WT and S400A/S422A double mutant were transiently transfected with or without OGT overexpression in HEK293 cells (Figure 7A) and SH-SY5Y neuroblastoma cells, respectively (Figure 7B). The cell viability was examined by MTS assay. Vectors-only or non-transfected cells were used as mock controls. The results showed that WT-Tau overexpression was toxic to both HEK293 and SH-SY5Y cells and the cell viability was significantly enhanced with OGT overexpression. In-contrast the double mutant showed no significant difference in cell viability with and without OGT overexpression. We further tested the neuronal toxicity of WT Tau and O-Tau fibrils and monomers to rat primary cortical neurons (Figure 7C). The samples at 40 nM were added to the neurons for 96 hr and the neurons were subjected to immunofluorescence imaging. Tau aggregates were stained with ThT. The results showed that WT Tau fibrils addition detected positive stain for ThT (green signal), severely affected neuronal morphology and resulted in loss of dendrites as shown by MAP2 staining (red). The mean neurite length was calculated using Image J analysis (Figure 7D). The result showed that there was a significant reduction in neurite length in the addition of WT Tau fibrils, but no such effect was observed in other conditions including WT Tau monomer, O-Tau monomers, and O-Tau fibrils addition.

**Figure 7.**
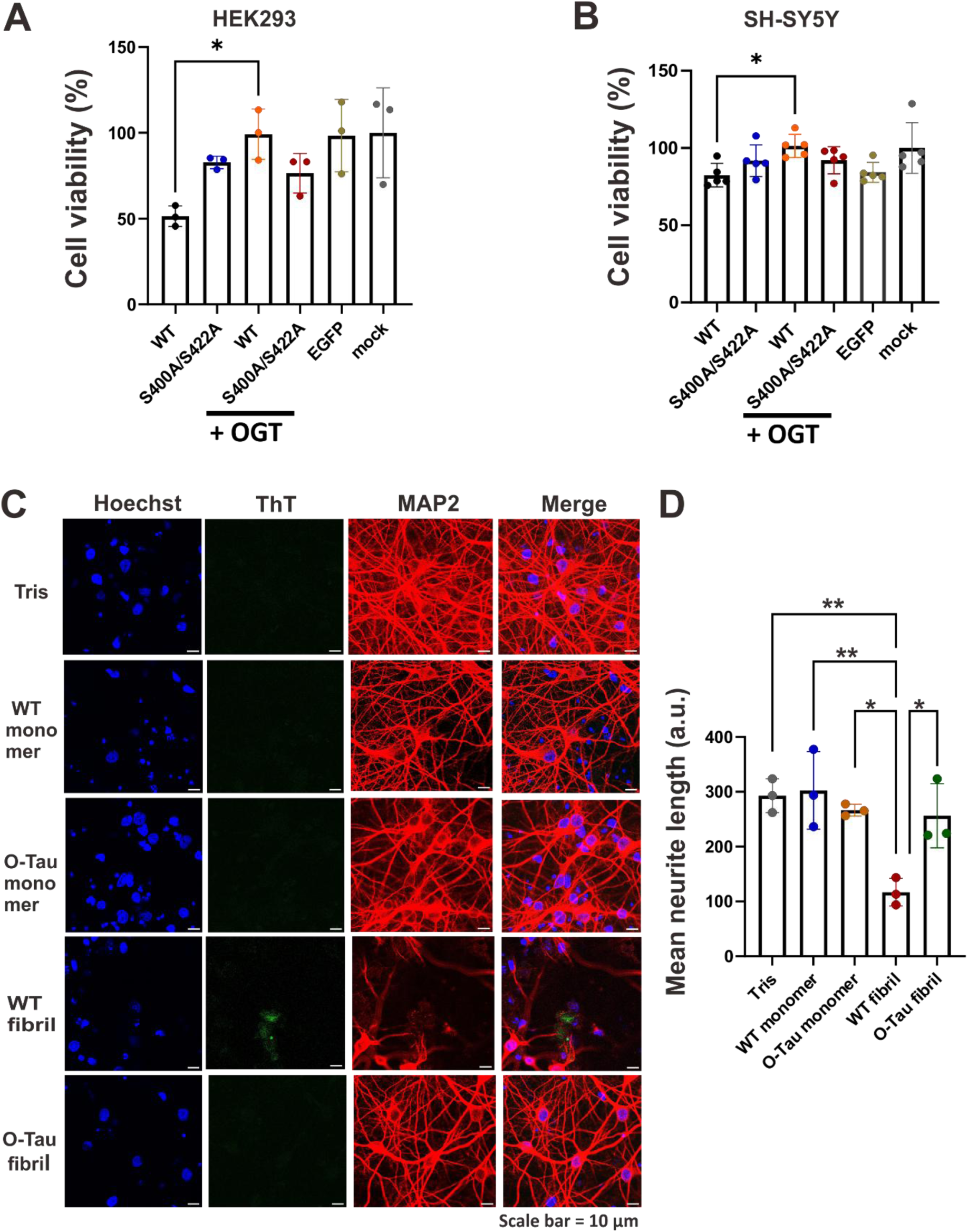
Tau toxicity analysis in HEK293 cells, SH-SY5Y cells, and rat primary cortical neurons. **(A-B)** Cell viability was assessed by MTS assay in (**A)** HEK293 cells **(B)** SH-SY5Y neuroblastoma cells expressing EGFP-Tau WT or the S400A/S422A double mut (mut), with or without OGT. Statistical significance was determined using one-way ANOVA, with Tukey’s multiple comparison post-test. \**P* < 0.05, \*\**P* < 0.01, \*\*\**P* < 0.001. **(C-D)** Representative immunofluorescence images of primary cortical neurons. Rat primary cortical neurons were treated with 40 nM of Tau-WT (WT) and O-Tau fibrils and monomers at 14 days after seeding, followed by 4 days of incubation. **(C)** Representative immunofluorescence images of primary cortical neurons. The experiment was performed for 3 independent repeats. **(D)** The mean neurite length was analyzed by ImageJ software. Statistical significance was determined using one-way ANOVA, with Tukey’s multiple comparison post hoc test. \**P* < 0.05, \*\**P* < 0.01.

### In-house novel *O*-GlcNAc-Tau-S422A antibody detects *O*-GlcNAcylation in vitro and in human brain samples, apart from highlighting the cross-talk relationship

A novel in-house monoclonal antibody, against Tau S422 *O*-GlcNAcylation was generated by differential screening for antibodies recognizing Tau-(412-430)-O-GlcNAc-S422 peptide. Monoclonal antibodies were generated by immunizing BALB/c mice with a KLH-conjugated glycopeptide (SSTGSIDMVDS(O-GlcNAc)PQLATLADCys), followed by hybridoma formation via splenocyte myeloma fusion. Antibody-producing clones were screened for site specific S422 *O*-GlcNAc modification, expanded, and purified from cell supernatants through Protein G affinity chromatography. This novel antibody could recognize S422 *O-*GlcNAcylation for recombinantly expressed O-Tau both in SP HP cation exchange chromatographic and *O*-GlcNAc enriched RP-HPLC fractions, where Tau-(412-430)-O-GlcNAc-S422 peptide, containing amino acid residues (412-430) with S422 site specific *O*-GlcNAc modification (Tau-O-S422) acted as a positive control, as shown in dot-blot analysis (Figure S7A). The antibody was also able to recognize S422 *O*-GlcNAcylation in WT Tau with OGT overexpression in HEK293 cells when immunoprecipitated against total Tau antibody Tau-5 (Figure S7B). Also, no S422-*O*-GlcNAcylation band was observed when recombinantly expressed *O*-GlcNAc-Tau was treated with GSK3ß kinase highlighting the prevalence of complex cross-talk relationships (Figure S7C). With the novel tool for recognizing S422 *O*-GlcNAcylation site, we used human brain tissues for further analysis. Next, immunohistochemical staining was subsequently conducted on amygdala and hippocampal regions of post mortem control and AD brain specimens. Representative images illustrate staining of control and AD tissues using the in-house *O*-GlcNAc-Tau-S422 antibody (Figure 8A) and with total Tau antibody Tau5^47^ (Figure 8B), respectively. The mean percentage of DAB-positive area per section was quantified across three independent control and three AD specimens. For each specimen, images from ten distinct regions within the amygdala and hippocampus were quantified using ImageJ, as shown in (Figure 8C, D). The results indicate a significant increase in total Tau levels in AD tissue, while the level of S422 *O*-GlcNAcylation compared to controls did not show statistical significance. However, the normalization of S422 *O*-GlcNAcylation to total Tau, showed a significantly higher relative *O*-GlcNAc level in control samples compared to AD (Figure 8E), suggesting that this modification is more abundant in the non-AD condition.

**Figure 8:**
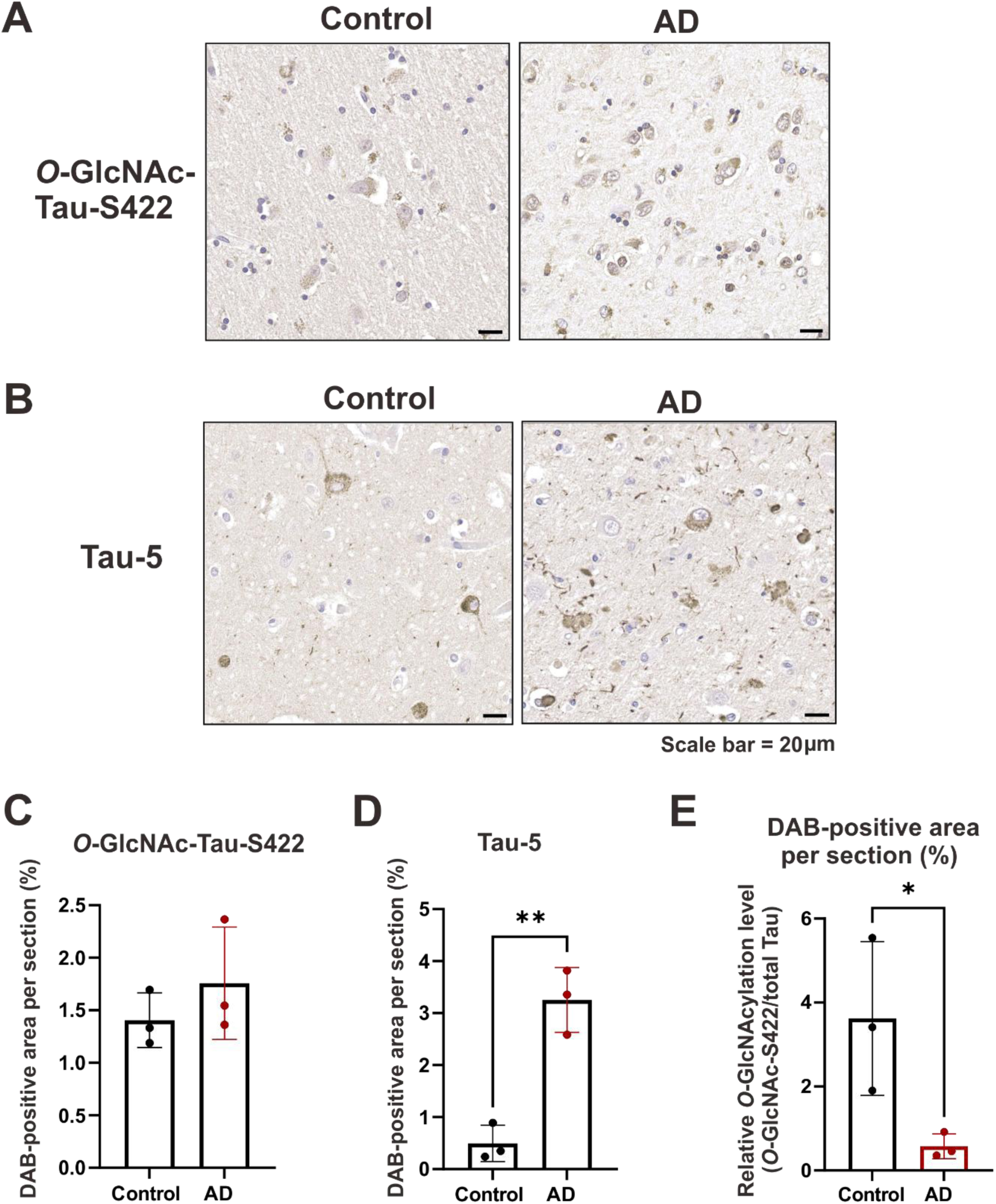
Immunohistochemical analysis of human brain tissue sections by in-house *O*-GlcNAc-Tau-S422 and total Tau Tau-5 antibodies. (A-B) Representative images of control and AD brain sections stained for (A) *O*-GlcNAc-Tau-S422 and **(B)** total Tau using Tau-5. Scale bar = 20 µm. (B) Representative images of control and AD brain tissue sections stained with the total Tau Tau-5 antibody. Scale bar = 20 µm. (C-D) Quantitative analysis of DAB-stained area in brain tissue sections from three control and three AD cases. The average percentage of DAB-stained area per section was calculated from 10 different regions for (C) *O*-GlcNAc-Tau-S422 staining and **(D)** Tau-5 staining. (E) *O*-GlcNAc-Tau-S422 signal normalized to total Tau level in control and AD brain samples. Statistical significance was determined using unpaired Student’s t-test. \**P* < 0.05, \*\**P* <0.01.

## Discussion

In our study we used various biophysical and biochemical techniques to study the detailed mechanism of Tau *O*-GlcNAcylation. We used a recombinant protein expression system in *E. coli* to overexpress *O*-GlcNAcylated-Tau-441 and validated the *O*-GlcNAc effect in vitro both globally and site-specifically. We detected several *O*-GlcNAc sites on recombinant full-length Tau-441 isoform, specifically located on the C-terminal domain. We found O-Tau aggregated slower with a more extended lag phase compared to WT. This effect may arise because C-terminal *O*-GlcNAcylation induces a more compact “paperclip-like” Tau conformation^48^, that may reduce aggregation propensity by enhancing solubility and structural stability. In seeding assays, O-Tau fibrils displayed reduced seeding capacity relative to WT fibrils. Addition of O-Tau seeds to WT monomers resulted in a shift from structured β-sheet–rich fibrils to more unstructured form, as indicated by an FTIR peak at ∼1637 cm⁻¹. Previous studies suggest *O*-GlcNAcylation enhances Tau solubility by destabilizing fibrillar structures and promoting soluble aggregate formation. Consistent with this, S400 *O*-GlcNAcylation has been shown to inhibit oligomerization and reduce aggregation and toxicity^23,49^. Our FCS data showed that O-Tau formed apparent heterogenous oligomeric assemblies of ∼4-5 Tau subunits, under LLPS condition compared to WT. *O*-GlcNAcylated Tau also exhibited enhanced FRET efficiency in cells, reflecting increased molecular proximity that is consistent with the oligomerization or conformational compaction. Together, these results support that O-Tau undergoes phase-separated clustering with tighter molecular interactions, linking reduced mobility indicated by (FCS) to increased nanoscale proximity shown by (FRET). Overall, *O*-GlcNAcylation appears to enhance solubility, extend early oligomerization, and delay fibril formation, thereby reducing seeding efficiency.

Among identified sites S400^25,50^ and S422, the latter emerged as a prominent and functionally relevant *O*-GlcNAc site. LC–MS/MS data repeatedly detected S422 modification, along with its previous pathological prevalence in AD brain^51^. S422 phosphorylation acted as one of the earliest biomarkers in preneurofibrillary tangles (NFT) in AD brain samples with low cognitive impairment^52^, where phosphorylation at S422 preceded conformational change and truncation in AD brain^53^. This suggests potential crosstalk between *O*-GlcNAcylation and phosphorylation at this site. The mutation of S422 alone and in combination with S400 markedly reduced overall *O*-GlcNAcylation reiterating its regulatory importance. The S400A/S422A double mutant also showed reduced phosphorylation at multiple C-terminal sites, suggesting that Tau regulation involves complex, context-dependent crosstalk rather than simple site competition between S/T modifications Depending on cellular conditions *O*-GlcNAcylation can differentially regulate phosphorylation at some sites^11^, including metabolic and oxidative stress^54^, and may induce allosteric effects that influence distant sites^39^. Hence, the balance between *O*-GlcNAcylation and phosphorylation as a dynamic sensor of cellular metabolic state, plays a crucial role in dictating the nature of the crosstalk.

For cellular transmission, we observed a significant increase in the transmission percentage of WT Tau upon OGT overexpression, and a progressive decrease from the single to the double mutants. This trend correlates with reduced phosphorylation in the C-terminal and microtubule-binding region (MTBR) in the S400A/S422A mutant. Phosphorylation of S400 and S422 were also detected in AD brains ^55–57^, with S422 site detected in extracellular vesicles from the AD brain tissues^58^. Phosphorylation at S422 impairs Tau transport^59^, and S400 affects its exosome associated secretion^60^. Tau can transmit via unconventional non-vesicular pathways^61^. The absence of these phosphorylation sites is likely to impact cellular propagation of Tau. The synergy between reduced *O*-GlcNAcylation and phosphorylation explains why these specific Tau mutants fail to transmit effectively. Previous studies suggest *O*-GlcNAcylation regulates protein trafficking and secretion via conventional and unconventional pathways^62^,with Tau release acting as a protective mechanism against intracellular accumulation^63^. FL Tau is present in CSF of AD patients, controls, and in transgenic mice before neuronal death, indicating active physiological release^64^. Structurally, pathological Tau forms β-sheet–rich, highly ordered fibrils associated with toxicity^65–67^. From our data, we hypothesize that *O*-GlcNAcylation promotes formation of smaller, less ordered, highly transmissible oligomeric assemblies that are less toxic than WT fibrils and may reduce toxic accumulation through enhanced transmission. Finally, identification of S422 as a major *O*-GlcNAc site led to generation of an in-house site-specific *O*-GlcNAc–Tau S422 antibody. This antibody specifically detected *O*-GlcNAcylated Tau at S422 in postmortem human brain tissue and revealed significantly reduced levels in AD compared to controls. This site-specific antibody may therefore serve as a tool to study S422 function and potentially act as a biomarker for AD, enabling further in vivo mechanistic and therapeutic investigations.

## Materials and Methods

All methods followed the standard guidelines from manufacturers. All culture reagents, unless specified, were purchased from Thermo Fisher Scientific (Waltham, MA, USA). All chemicals, unless specified, were obtained from Sigma-Aldrich (St. Louis, MO, USA).

### Expression and purification of WT-Tau and *O-*GlcNAc-Tau

Full-length Tau-441 (WT) in pET29b vector was transformed in *E. coli* BL21 (DE3). A single colony was picked up and cultured in 2L LB broth with 50 ug/mL kanamycin at 37°C. The culture was shaking at 200 rpm until OD_600_ reaching 0.8 and followed by protein induction by 1 mM IPTG at 37°C for 3 hr. Bacterial cells were then pelleted down at 7000 g for 15 min at 4°C. The pellet was resuspended in lysis buffer containing 20 mM MES, pH 6.5, 1 mM MgCl_2,_ 100 mM NaCl, 5 mM DTT and 2 mM EDTA in the presence of protease inhibitor cocktail III (Millipore) and lysed by microfluidizer for 30 cycles and boiled at 80°C for 30 min with occasional stirring. The cell lysate was then centrifuged at 20,000 g for 45 min at 4°C to generate supernatant which was further purified via cation exchange column HiTrap SP Sepharose HP (GE healthcare biosciences) on FPLC system. The sample was loaded using Buffer A (20 mM MES, pH 6.5, 1 mM MgCl_2,_ 5 mM NaCl, 5 mM DTT, 2 mM EDTA). Before sample loading, the column was first equilibrated by 30% buffer B (20 mM MES, pH 6.5, 1 mM MgCl_2,_ 500 mM NaCl, 5 mM DTT, 2 mM EDTA), washed with 30 % buffer B. A gradient of 30% to 100% was used to elute Tau using elution buffer B. The fractions were collected and dialyzed overnight in presence of 1 mM DDT with dialysis buffer having similar composition to Buffer A. The protein was next purified by heparin HP column (GE healthcare biosciences) pre-equilibrated by 40% Buffer B. Tau fractions were eluted and collected in a gradient from 40% to 80% Buffer B and dialyzed against 20 mM Tris-HCl, pH 7.4, overnight followed by 4 hr at 4°C and Snap frozen in Liquid N_2_ and kept in −20°C for long term storage.

For *O*-GlcNAc-Tau (O-Tau) purification, Tau-441 (WT) in pET29b vector and OGT were co-transformed in *E. coli* BL21 (DE3). A single colony was picked up and cultured in 2L of LB broth with 100 µg/mL ampicillin and 25 ug/mL kanamycin respectively. The culture was shaken at 200 rpm until OD_600_ reaching 0.8 and followed by induction by 1 mM IPTG at 37°C for 3 hr. Bacterial cells were then pelleted down at 7,000 g for 15 min at 4°C. The pellet was resuspended in lysis buffer containing 20 mM MES, pH 6.5, 1 mM MgCl_2,_ 100 mM NaCl, 5 mM DTT, and 2 mM EDTA in the presence of protease inhibitor cocktail III (Millipore) and lysed by microfluidizer for 30 cycles and boiled at 80°C for 30 min with occasional stirring. The cell lysate was then centrifuged at 20,000 g for 45 min at 4°C to generate supernatant which was further purified via cation exchange column HiTrap SP Sepharose HP (GE healthcare biosciences) on FPLC system. The sample was loaded using Buffer A (20 mM MES, pH 6.5, 1 mM MgCl_2,_ 5 mM NaCl, 5 mM DTT, 2 mM EDTA). Before sample loading, the column was first equilibrated by 30% buffer B (20 mM MES, pH 6.5, 1 mM MgCl_2,_ 500 mM NaCl, 5 mM DTT, 2 mM EDTA), washed with 30 % buffer B. A gradient of 30% to 100% was used to elute Tau using elution buffer B. The eluted protein was dialyzed overnight at 4°C in 20 mM Tris-HCl pH 7.4. *O-*GlcNAc enrichment was further performed by Reverse Phase-High Performance Liquid Chromatography (RP-HPLC) using a Semi-preparative Zorbax 300SB C8 column (Agilent). The column was first equilibrated by 20% of buffer B (90 % ACN, 0.1% v/v of TFA at pH 2-2.5). At a time 250 ug of Tau was injected into the HPLC machine. The protein was eluted between 20-30% of buffer B. The different elution peaks obtained were collected and lyophilized to dryness followed by Tau-*O*-GlcNAc confirmation by Dot blot and Mass spectrometric analysis.

### Dot blot Analysis

Two µL each of WT-Tau, O-Tau SP (*O*-GlcNAc Tau fraction purified by SP Sepharose), Tris buffer and the RP-HPLC fractions were dotted on a nitrocellulose membrane, dried and blocked with 1X carbon-free blocking buffer (Vector laboratories) for 1 hr. The membrane was then probed with primary antibodies anti-*O-*GlcNAc-TauS400 antibody (1:500, Anaspec) RL2 (1:500, Invitrogen), and Tau-5 (1:5000, Invitrogen) followed by washing with TBST. The membrane was then blotted with secondary antibodies namely, anti-rabbit (1:500, GeneTex), anti-mouse (1:500, GeneTex), and anti-mouse (1:5000), respectively and was finally detected by chemiluminescence using ECL substrate kit (Merck Millipore).

### Mass spectroscopic analysis

Molecular mass of the protein in the RP-HPLC fractions were determined by LC-MS experiment on a SELECT SERIES Cyclic IMS (Waters, Wilmslow, U.K.), equipped with Waters ACQUITY UPLC I-Class (Waters, Milford, MA).

MALDI-TOF measurements were performed on RP-HPLC fractions by an Ultraflex II MALDI-TOF/TOF mass spectrometer (Bruker Daltonik GmbH, Bremen, Germany). Mass spectra were obtained in the range of mass to charge ratio (*m*/*z*) from 10,000 to 100,000 with linear mode.

The *O*-GlcNAc-Tau protein was in-solution digested by trypsin at trypsin-to-protein ratio 1:50, and *O*-GlcNAcylation sites were detected by LC-MS/MS on an Orbitrap Fusion mass spectrometer (Thermo Fisher Scientific, San Jose, CA) equipped with EASY-nLC 1200 system (Thermo, San Jose, CA, US) and EASY-spray source (Thermo, San Jose, CA, US), and analyzed by MASCOT software. The experiments were performed by GRC Mass Core Facility of Genomics Research Center, Academia Sinica, Taipei, Taiwan.

### ThT assay

WT and *O*-GlcNAc-Tau monomers at 5 µM were incubated in 20 mM Tris buffer, pH 7.4, in the presence of 1.25 µM heparin at ratio 4:1 and 10 µM ThT, at 25°C. The mixture was aliquoted at 40 µL volume in a 384 well plate and fluorescence intensity was measured over a period of time at fixed intervals, along with occasional shaking, using a SpectraMax M5 microplate reader (excitation at 442 nm; emission at 485 nm).

### Preparation of Tau fibrils

WT and *O*-GlcNAc-Tau at 5 µM in 20 mM Tris buffer, pH 7.4, were fibrillized in the presence of 1.25 µM heparin and 10 µM ThT over a period of time to form fibrils. The preformed tau fibrils were sonicated using Ultrasonic Cleaner 3510 (Branson) for 20 min prior to the experiments.

### ThT Seeding Assay

WT and *O*-GlcNAc-Tau monomers at 5 µM were centrifuged at 17,000g for 30 min, then incubated in 20 mM Tris buffer, pH 7.4, in the presence of 5% v/v of WT and *O-*GlcNAc-Tau preformed seeds, respectively along with 1.25 µM heparin and 10 µM ThT, at 25°C. The ThT fluorescence intensity was monitored for 72 hr with 30 sec agitations per hr, and the fluorescence intensity was detected at fixed intervals using a SpectraMax M5 microplate reader (excitation at 442 nm; emission at 485 nm). The end product of the seeding assay was collected and stored at 4°C for further analysis.

### Transmission electron microscopy

To determine the morphology of WT and *O*-GlcNAc-Tau aggregates, 10 µL of the final products at the end of ThT aggregation and seeding assay were applied onto 400-mesh Formvar carbon-coated copper grids (EMS Inc., Hatfield, PA, USA) for 5 min, rinsed, and then negatively stained by 2 % PTA. A Hitachi H-7000 transmission electron microscope (TEM) (Hitachi Inc., Tokyo, Japan) with an accelerating voltage of 120 kV was employed to examine the samples.

### Fourier-transform infrared spectroscopy

To determine conformation of WT and *O*-GlcNAc-Tau aggregates, 10 µL of the final products at the end of ThT aggregation and seeding assay were analyzed by ATR-Fourier-transform infrared microscopy (FTIR). The samples were centrifuged at 17,000 g for 30 min at 4°C and washed with deionized water, then resuspended into D_2_O before applied onto PIKE Miracle ATR sampling accessory and dried with cool air. Data between 1,600 and 1,700 cm^-1^ were collected and normalized.

### Liquid-liquid phase separation and FRAP analysis

Tau at 2 mg/mL was labelled with fluorescent dye Alexa fluor 488, at a protein to dye ratio 10:1, in 10 mM HEPES buffer pH 7.4 overnight at 4°C and excess dye was filtered using gel filtration column (Zeba spin desalting columns, microspin G25). The final concentration and degree of labelling was calculated using a nanodrop. In vitro LLPS was studied for 10 µM WT-Tau and *O*-GlcNAc-Tau with increasing concentration of NaCl (50 mM,100 mM, 150 mM) in 0.1 mM EDTA, 2 mM DTT in presence of crowding agent 10% PEG 8000. Spherical droplet formation was observed under SP8X Confocal Super resolution microscope. Mean droplet area and diameter were calculated by Image J software. Droplet dynamics for WT and O-Tau was studied using fluorescence recovery after photobleaching (FRAP) analysis. 10 µM of Alexa flour 488 labelled WT and O-Tau in HEPES buffer, pH 7.4, 50 mM NaCl and 10 % PEG 8000 was dropped on glass bottomed chamber slides at 100 X magnification (oil lens) on a Leica Hy Volution SP8 confocal microscope. Droplets were imaged with a 488 nm laser and 450/50 nm detection filters. A small region (ROI = 0.3 µm) was chosen and bleached with 3 frames of full laser power. Post bleaching fluorescence recovery was detected up to 270 sec.

### Fluorescence correlation spectroscopy (FCS

Ten µM of WT and O-Tau monomers were doped with 50 nM of Alexa Flour 488 labelled WT and O-Tau under LLPS conditions (10 mM HEPES buffer, pH 7.4, 50 mM NaCl, 10 % PEG 8000), FCS measurements were acquired using time resolved fluorescence microscope (MicroTime 200, PicoQuant GmbH), built on a modified Olympus IX71 inverted microscope, equipped with a 100x oil-immersion objective and a pulsed 488 nm excitation laser. Fluorescence signals were detected using single photon avalanche diode (SPAD) photon counting detectors. Fluorescence intensity fluctuations were recorded for 180 s and analyzed using SymphoTime software. Autocorrelation curves were generated from fluorescence intensity fluctuations and fitted using a three-dimensional Brownian diffusion model incorporating triplet-state relaxation and two-component diffusion. The correlation function was described as follows:

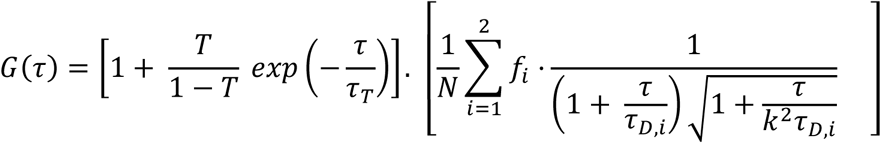

where, N is the average number of fluorescent particles within the confocal volume, *f*_*i*_ represents the fractional contributions of each diffusing species (∑ *f*_*i*_ = 1), τ_*D,i*_ represents diffusion time of each component, and *k* = *z*_0_/ω_0_ is the structure parameter relating the axial and lateral dimensions of the confocal volume. The first bracket accounts for the triplet state dynamics, with T and τ_*T*_ represent triplet state fraction and triplet relaxation lifetime, respectively. Diffusion coefficients were obtained using *D_i_* = ω^2^_0_/(4τ_D,i_), where ω_0_ was calibrated using Alexa Flour 488 with a known diffusion coefficient^68^.Hydrodynamic radii of O-Tau species within condensates were estimated using the Stokes-Einstein equation.

### Fluorescence lifetime imaging microscopy-Förster resonance energy transfer (FLIM-FRET) analysis

FRET experiments were conducted in HEK293 cells. Plasmids encoding EGFP and mCherry vectors only (control), EGFP-tagged WT or S400A/S422A (donors) together with mCherry tagged WT Tau or S400A/S422A (acceptors) double mutant (acceptors), were co-transfected with or without OGT overexpression at equimolar plasmid ratios to ensure balanced donor and acceptor expression. FRET measurements were acquired at 48 hours after transfection using a time resolved fluorescence microscope (MicroTime 200, PicoQuant GmbH) built on a modified Olympus IX71 inverted microscope platform. Donor fluorescence lifetimes were recorded in the absence or presence of acceptor constructs using time-correlated single-photon counting (TCSPC) detection. Acquisition settings were optimized to achieve sufficient photon statistics while minimizing photobleaching. Regions of interest were selected within the cytoplasm of transfected cells while excluding saturated or low-intensity regions. FRET efficiency (E) was calculated from donor lifetime reduction using the equation, *E* = 1 − (τ _DA_ / τ_D_), where τ _DA_ represents the donor fluorescence lifetime in the presence of acceptor and τ_D_ corresponds to the donor-only lifetime controls. Background correction was applied prior to analysis. Fluorescence lifetime fitting and FRET analysis were performed using SymPhoTime software (PicoQuant) following standard TCSPC-based FLIM analysis workflows.

### *O*-GlcNAc-Tau mutants’ generation by site-directed mutagenesis

EGFP-Tau-441 and mCherry-Tau-441 plasmids were previously constructed in the lab^24^. *O*-GlcNAc-Tau single and double mutations at S400 and S422 sites to alanine was performed by site-directed mutagenesis on EGFP-Tau-441 and mCherry-Tau-441 plasmids, respectively, by Site Directed Mutagenesis Polymerase chain reaction using phosphorylated forward and reverse primer, in the presence of KAPAHiFi ^TM^ DNA polymerase. The base alteration was verified by sequencing. The forward and reverse primer sequences for S400A were 5’ AAGTCGCCAGTGGTGGCTGGGGACACGTCTCCA 3’ and 5’ TGGAGACGTGTCCCCAGCCACCACTGGCGACTT 3’, respectively. The forward and reverse primer sequences for S422A were 5’ CGACATGGTAGACGCGCCCCAGCTCGCCACG 3’ and 5’ CGTGGCGAGCTGGGGCGCGTCTACCATGTCG 3’, respectively. Various single and double EGFP and mCherry Tau mutants were generated: EGFP-Tau-441-S400A (gT-S400A), EGFP-Tau-441-S422A (gT-S422A), p-EGFP-Tau-441-S400A_S422A (gT-S400A/S422A), mCherry-Tau-441-S400A (mT-S400A), mCherry-Tau-441-S422A (mT-S422A) and mCherry-Tau441-S400A_S422A (mT-S400A/S422A).

### Cell culture

HEK293 (Human embryonic kidney 293 cells) cells were cultured in DMEM (Dulbecco’s Modified Eagle Medium) supplemented by 10% (v/v) FBS (Fetal Bovine Serum) and 1% penicillin/streptomycin. Similarly, human neuroblastoma SH-SY5Y were cultured in DMEM medium, and supplemented by 10% FBS, 1% penicillin/streptomycin. GFP expressing HEK293 stable cell line was cultured in DMEM medium, and supplemented by 10% FBS, 1% penicillin/streptomycin, along with 0.5 mg/mL G418 marker. Cells were maintained in 5% CO_2_ under humidified conditions at 37°C incubator and were passaged when 80-90% confluence was reached.

### Immunoprecipitation and western blot analysis

The plasmids gT-WT, gT-S400A, gT-S422A and gT-S400A/S422A were transiently transfected with Lipofectamine 3000 in the presence of OGT overexpression at (1:3) for 3 days. Cells were lysed in Triton-X lysis buffer containing 500 mM Tris-HCl, pH 8.0, 137 mM NaCl, 1% Triton X-100, 2 mM EDTA, OGA inhibitor (Thiamet-G), N-acetylhexosaminidase inhibitor (PUGNAc) and protease inhibitors. The supernatant was incubated in magnetic beads coated with total Tau antibody Tau-5, at 4°C overnight. Magnetic beads were washed five times with wash buffer (10 mM Tris pH 7.4, 1 mM EDTA, 1 mM EGTA pH 8.0, 150 mM NaCl, 1% Triton X-100, 1 µM TMG, PUGNAc and protease inhibitor), and the proteins were eluted by 2X SDS sample buffer. The eluted protein was run through SDS-PAGE followed by transfer to PVDF membrane. The membrane was blocked with 1X Carbon-free blocking buffer followed by probing with RL2 (1:500), OGT (1:1000), and Tau-5 (1:5000) primary antibodies respectively. The membrane was then washed with TBST buffer and probed with anti-mouse (1:500), anti-rabbit (1:5000), and anti-mouse (1:5000) secondary antibodies respectively. The membrane was detected by chemiluminescence using ECL substrate kit. The blots were quantified using Image J analysis.

### In vitro Tau phosphorylation

In vitro Tau phosphorylation was performed following a similar protocol from previous literature^43^. In our case, recombinantly expressed WT and O-Tau were incubated with GSK-*3ß* at 50:1 ratio in the presence of 2 mM ATP, 5 mM EGTA, 5 mM MgCl_2_, and 2 mM DTT in 40 mM HEPES buffer at pH 7.4 for 12 hr at 30 °C with continuous shaking at 300 rpm in an Eppendorf thermomixer. After the completion of the phosphorylation reactions, the samples were boiled at 98°C in order to precipitate kinases, then centrifuged at 20,000 g for 20 min and the supernatant was carefully collected and stored.

### Detection of in vitro cross-talk relationship between *O*-GlcNAcylation and phosphorylation in Tau

In order to address the cross-talk relationship at S396 and S404 sites at the vicinity of S400 *O*-GlcNAc site, Tris, recombinant WT and O-Tau with or without GSK-*3ß* phosphorylation were dotted on NC membrane and the previously mentioned protocol of dot blot analysis was performed using anti *O*-GlcNAc detecting antibodies namely *O*-GlcNAc-Tau-S400 (1:500) and RL2 (1:500) and site-specific Tau phosphorylation detecting antibodies namely S396 (1:2500) and S404 (1:2500) respectively. Similarly, in HEK293 cells, gT-WT and gT-S400A/S422A plasmids with or without OGT overexpression and EGFP were transiently transfected using Lipofectamine 3000 and phosphorylation and total tau expressions at sites S396 (1:2500) and S404 (1:2500) sites, respectively, were detected by Western blot analysis following the previously mentioned protocol and quantified by Image J analysis.

### Cell-to-cell transmission analysis using Flow cytometry

Cellular transmission was studied in HEK293 cells, where cells were transiently transfected using Lipofectamine 3000 with EGFP and mCherry only and EGFP and mCherry-Tau WT, S400A, S422A and S400A/S422A mutants respectively, with or without OGT overexpression and TMG administration at 1 µM. The protein was over expressed for 72 hr after which equal volumes of EGFP-Tau and mCherry Tau expressing cells were co-cultured at 300,000 cells/well in 6 well plates. EGFP and mCherry alone as co-culture were used as control. The co-cultured cells were harvested after 48 hr. A FACS Aria II flow cytometer (BD biosciences) equipped with 488 nm and 561 nm lasers having GFP and Yellow-PE-Texas red filters was used. Dual positive cells out of all fluorescent were quantified for 10,000 events per sample. The transmission was indicated by percentage of cells with dual fluorescence in all fluorescent cells. The results were analyzed by FlowJo software (FlowJo LLC, US). The experiments were performed for three replicates.

### Primary cortical neuron extraction and culture

Primary cortical neurons were isolated from E17 rat embryos, following the National Institute of Health Guidelines for Animal Research and Taiwan Animal Protection Law, approved by Academia Sinica Institutional Animal Care and Use Committee (IACUC 22-05-1853). Pregnant Sprague-Dawley rats carrying E17 embryos were obtained from BioLASCO Taiwan Co., Ltd. Neurobasal medium (Cat. No.: 21103-049) was supplemented with B27 supplement (Cat. No.: 17504044), 0.25% GlutaMAX-I (Cat. No.:35050-061) and Penicillin/ Streptomycin (Cat. No.: 15140122). The plates were coated with 25 μg/ml poly-D-lysine hydrobromide (Cat. No.: P6407) and washed with autoclaved sterile double distilled water twice. Pregnant Sprague-Dawley rats were euthanized by CO_2_ inhalation for 5 minutes. The uterus was taken out and placed on a new petri dish on ice and the following procedures were performed in the BSC. Fetuses were taken out without directly touching the brain followed by placing them on a new petri dish on ice. A Y shaped incision was made carefully on the skull to expose the whole brain. The hole brain was removed and kept on ice in a serum-free DMEM medium containing 1,000 units/mL of penicillin and 1,000 μg/mL of streptomycin, on ice. The meninges, blood vessels and non-cortical brain regions were carefully removed to isolate the cortex. The cortex was carefully transferred from DMEM to to ∼5 mL 0.25% trypsin and incubated in 37°C water bath for 15 min, followed by centrifugation at 1000 rpm for 5 min and trypsin was removed. The pellet was resuspended in 5 mL of 1 mg/mL trypsin inhibitor (Cat. No.: 17075029), and gently vortexed for 10 times, followed by centrifugation at 1000 rpm for 5 min to remove trypsin inhibitor. The pellets were further resuspended in pre-warmed neurobasal medium and filtered through a 40 μm cell strainer. The cell suspension was centrifuged at 1000 rpm for 5 min and the pellet was re-suspended in pre-warmed neurobasal medium. The medium was changed after 30 min of cell seeding to remove debris and non-adherent cells. Half of the of medium was changed every 3 days.

### Immunofluorescence

The primary cortical neurons were grown for 14 days on coverslips coated with poly-D-lysine until maturity and were treated with 20 mM Tris buffer as negative control and extracellular pre-formed WT and O-Tau monomer and fibrils respectively at 40 nM concentration. After treatment for 4 days, neurons were washed with PBS buffer and fixed with 4 % paraformaldehyde for 15 min at room temperature. After washing with 1% PBST thrice, cells were permeabilized with 0.5% Triton X-100 in PBS for 10 min. After washing with 1% PBST 3 times, cells were next blocked with 1% BSA in PBST followed by primary antibody staining with anti-guinea pig MAP2 (1:1000) overnight at 4°C. The neurons were washed with 1% PBST thrice followed by subsequent incubation with 10 µM of ThT, Hoechst 33358 (10 µg/mL) and secondary antibody anti-guinea pig Alexa^594^ (1:500) for 30 min. The cells were washed first with 1% PBST for 3 times then washed with 1X PBS. The cells were mounted by 10 mM Tris in 50 % glycerol mounting solution and imaged by SP-8 confocal microscopy using 63X magnification. Data was analyzed by LAS-X software. Mean neurite length was quantified for 3 independent repeats by Image J analysis using NeuronJ plugin.

### Cell-cytotoxicity analysis

The plasmids mCherry-Tau WT and S400A/S422A were transiently transfected in HEK293 cells and SH-SY5Y cells with or without OGT overexpression, respectively. Vector alone was transfected as control. Proteins were overexpressed for 72 hr and then seeded in 96 well plate for 24 hr, along with cells only as control, respectively. Next MTS reagent was added and cells were incubated for 2 hr after which the absorbance was measured at 490 nm and the percent cell viability was quantified.

### Antibody Production

Tau-(412-430) peptide and Tau-(412-430)-O-GlcNAc-S422 peptide, which is the peptide containing amino acid residues (412-430) with or without S422 site specific *O*-GlcNAc modification, respectively, were chemically synthesized by the Genomics Research Center Peptide Core, Academia Sinica. Monoclonal antibodies were produced by LTK BioLaboratories, Taiwan using the peptide (SSTGSIDMVDS(O-GlcNAc)PQLATLADCys) corresponding to the epitope region of Tau as the antigen containing the site specific S422-*O*-GlcNAc modification, Hybridoma cell lines were generated through immunization of BALB/c mice with the peptide conjugated to keyhole limpet hemocyanin (KLH), followed by fusion of splenocytes with myeloma cells. Clones secreting specific antibodies were selected by enzyme-linked immunosorbent assay (ELISA) and validated against recombinant Tau-441. Hybridoma cells were subsequently expanded, and monoclonal antibodies were purified from culture supernatants using Protein G affinity chromatography.

### Immunohistochemical staining of human brain tissues

Human tissue related usage and procedures were approved by Human Subjects Research Ethics, Academia Sinica, Taiwan. Post-mortem human brain tissues were obtained from Dr. Lee-Way Jin’s laboratory at The Alzheimer’s Disease Center, University of California, Davis Medical Center, Sacramento, CA, 95817, USA. The study was performed under the IRB of Academia Sinica (AS-IRB-BM-14012). Paraffin-embedded blocks of 5µm thickness of anterior hippocampus and amygdala specimens of human controls aged 75 (F, PMI 3.5), 80 (M, PMI 18.1), 84 (F, PMI 10.9) with Braak & Braak stages I, 0, I respectively and of high Alzheimer’s disease aged 85 (F, PMI 4), 75 (F, PMI 3), 81 (M, PMI 4) with Braak & Braak stages V brain sections, respectively, were stained using a commercially available detection system (Bio SB). Deparaffinization with xylene, of the tissue sections were performed with a GRC(8)–Leica Biosystems Autostainer XL platform. Deparaffinization was followed by rehydration by washing through graded ethanol in water solvent ending in pure ddH_2_ O. Sections were maintained in phosphate - buffered saline (PBS) at 4 °C. Antigen retrieval was achieved by heat-induced treatment in citrate buffer (10 mM citric acid, pH 6.0, 0.05% Tween-20). Slides were placed in preheated retrieval solution and maintained at 90–95 °C for 20 min under controlled conditions. Samples were then allowed to equilibrate gradually to ambient temperature. After cooling, sections were rinsed for 5 min in PBST (low-salt PBS containing 0.1% Triton X-100). Endogenous peroxidase activity was suppressed by incubation with hydrogen peroxide reagent supplied in the kit for 5 min at room temperature, followed by a 5-min PBST rinse. Non-specific binding was minimized using 3% bovine serum albumin (BSA) prepared in PBST (PBS supplemented with 0.3% Triton X-100) for 30 min at ambient temperature. For *O*-GlcNAc–targeted staining, an additional blocking step was included using 1% BSA prepared in carbon-free buffer (Vector Laboratories) containing 0.05% Tween-20 for 30 min at room temperature. Sections were subsequently rinsed with PBST. A hydrophobic barrier was applied around each specimen prior to primary antibody exposure. Samples were incubated overnight at 4 °C with (in-house) anti–O-GlcNAc–Tau-S422 antibody (O-Tau-S422; 1:200 dilution) or total Tau antibody Tau-5 (1:500 dilution). Following primary incubation, slides were subjected to three PBST washes (5–10 min each). A linker/enhancer solution (Bio SB) was applied for 30 min at room temperature, followed by three additional PBST washes. Polymer-HRP conjugate was then applied for 30 min at ambient temperature, followed by five PBST rinses. Chromogenic detection was carried out using 3,3 ′-diaminobenzidine (DAB) treatment. Tissue sections were exposed to substrate for 30-40 sec while monitoring signal development microscopically. The reaction was terminated by immediate immersion in PBS. Slides were rinsed, dehydrated through graded ethanol series, cleared, and mounted using the GRC (8)–Leica Autostainer XL system. Digital images were acquired and analyzed using Aperio ImageScope software.

## Statistical Analysis

All experiments were performed in at least triplicates for individual sample. Statistical analysis was performed using GraphPad Prism (GraphPad Software, US) as the means ± standard deviations (SD). Student’s t-test and ANOVA were used for studying significance at * P < 0.05, ** P < 0.01, and *** P < 0.001.

## Acknowledgments

We thank the scientific funding from the Ministry of Science and Technology, Taiwan (MOST), and Academia Sinica (AS), Taiwan (AS-GC-108-07, AS-GC-111-L03, AS-SUMMIT-109, AS-KPQ-109-BioMed, 104-2320-B-001-013-MY3, MOST 106-0210-01-15-02, MOST 107-0210-01-19-01, MOST-108-3114-Y-001-002). We thank Mr. Chein-Hung Chen of the mass spectrometry core, the Peptide synthesis Core, the Imaging Core Facility, Flow Cytometry Facility at the Genomics Research Center, AS, and TEM core facility, AS, for providing technical support.

## Author contributions

D.K. designed and performed all biochemical and cellular experiments, analyzed the data and wrote the manuscript. W.W.C. processed the human postmortem tissue samples and performed the initial immunohistochemistry experiment; repeat experiments were subsequently performed by D.K. W.C.L. performed primary cortical neuron extraction and culture. L.W.J. provided human brain samples. W.C.H. assisted with FCS measurements and data analysis. Y.R.C. conducted the research direction and edit the manuscript. All authors reviewed and approved the manuscript.

## References

1 Silva, M. V. F. et al. Alzheimer’s disease: risk factors and potentially protective measures. J Biomed Sci 26, 33 (2019). 10.1186/s12929-019-0524-y

2 DeTure, M. A. & Dickson, D. W. The neuropathological diagnosis of Alzheimer’s disease. Mol Neurodegener 14, 32 (2019). 10.1186/s13024-019-0333-5

3 Braak, H. & Braak, E. Neuropathological stageing of Alzheimer-related changes. Acta neuropathologica 82, 239–259 (1991).

4 Zhang, H., Cao, Y., Ma, L., Wei, Y. & Li, H. Possible Mechanisms of Tau Spread and Toxicity in Alzheimer’s Disease. Front Cell Dev Biol 9, 707268 (2021). 10.3389/fcell.2021.707268

5 Buchholz, S. & Zempel, H. The six brain-specific TAU isoforms and their role in Alzheimer’s disease and related neurodegenerative dementia syndromes. Alzheimer’s & dementia: the journal of the Alzheimer’s Association 20, 3606–3628 (2024). 10.1002/alz.13784

6 Guo, J. L. & Lee, V. M. Seeding of normal Tau by pathological Tau conformers drives pathogenesis of Alzheimer-like tangles. J Biol Chem 286, 15317–15331 (2011). 10.1074/jbc.M110.209296

7 Kaufman, S. K. et al. Tau Prion Strains Dictate Patterns of Cell Pathology, Progression Rate, and Regional Vulnerability In Vivo. Neuron 92, 796–812 (2016). 10.1016/j.neuron.2016.09.055

8 Amorim, I. S. et al. A seeding-based neuronal model of tau aggregation for use in drug discovery. PloS one 18, e0283941 (2023). 10.1371/journal.pone.0283941

9 Alquezar, C., Arya, S. & Kao, A. W. Tau Post-translational Modifications: Dynamic Transformers of Tau Function, Degradation, and Aggregation. Front Neurol 11, 595532 (2020). 10.3389/fneur.2020.595532

10 Xia, Y., Prokop, S. & Giasson, B. I. “Don’t Phos Over Tau”: recent developments in clinical biomarkers and therapies targeting tau phosphorylation in Alzheimer’s disease and other tauopathies. Mol Neurodegener 16, 37 (2021). 10.1186/s13024-021-00460-5

11 Liu, F., Iqbal, K., Grundke-Iqbal, I., Hart, G. W. & Gong, C. X. O-GlcNAcylation regulates phosphorylation of tau: a mechanism involved in Alzheimer’s disease. Proc Natl Acad Sci U S A 101, 10804–10809 (2004). 10.1073/pnas.0400348101

12 Ryan, P. et al. O-GlcNAc Modification Protects against Protein Misfolding and Aggregation in Neurodegenerative Disease. ACS chemical neuroscience 10, 2209–2221 (2019). 10.1021/acschemneuro.9b00143

13 Marotta, N. P. et al. O-GlcNAc modification blocks the aggregation and toxicity of the protein α-synuclein associated with Parkinson’s disease. Nat Chem 7, 913–920 (2015). 10.1038/nchem.2361

14 Balana, A. T. et al. O-GlcNAc forces an α-synuclein amyloid strain with notably diminished seeding and pathology. Nat Chem Biol 20, 646–655 (2024). 10.1038/s41589-024-01551-2

15 Gong, C. X., Liu, F. & Iqbal, K. O-GlcNAcylation: A regulator of tau pathology and neurodegeneration. Alzheimers Dement 12, 1078–1089 (2016). 10.1016/j.jalz.2016.02.011

16 Hardivillé, S. & Hart, G. W. Nutrient regulation of signaling, transcription, and cell physiology by O-GlcNAcylation. Cell Metab 20, 208–213 (2014). 10.1016/j.cmet.2014.07.014

17 Yang, X. & Qian, K. Protein O-GlcNAcylation: emerging mechanisms and functions. Nat Rev Mol Cell Biol 18, 452–465 (2017). 10.1038/nrm.2017.22

18 Lefebvre, T. et al. Evidence of a balance between phosphorylation and O-GlcNAc glycosylation of Tau proteins--a role in nuclear localization. Biochim Biophys Acta 1619, 167–176 (2003). 10.1016/s0304-4165(02)00477-4

19 Liu, F. et al. Reduced O-GlcNAcylation links lower brain glucose metabolism and tau pathology in Alzheimer’s disease. Brain 132, 1820–1832 (2009). 10.1093/brain/awp099

20 Graham, D. L. et al. Increased O-GlcNAcylation reduces pathological tau without affecting its normal phosphorylation in a mouse model of tauopathy. Neuropharmacology 79, 307–313 (2014). 10.1016/j.neuropharm.2013.11.025

21 Borghgraef, P. et al. Increasing brain protein O-GlcNAc-ylation mitigates breathing defects and mortality of Tau.P301L mice. PLoS One 8, e84442 (2013). 10.1371/journal.pone.0084442

22 Hastings, N. B. et al. Inhibition of O-GlcNAcase leads to elevation of O-GlcNAc tau and reduction of tauopathy and cerebrospinal fluid tau in rTg4510 mice. Mol Neurodegener 12, 39 (2017). 10.1186/s13024-017-0181-0

23 Yuzwa, S. A., Cheung, A. H., Okon, M., McIntosh, L. P. & Vocadlo, D. J. O-GlcNAc modification of tau directly inhibits its aggregation without perturbing the conformational properties of tau monomers. J Mol Biol 426, 1736–1752 (2014). 10.1016/j.jmb.2014.01.004

24 Chen, G. J. et al. Tau destabilization in a familial deletion mutant K280 accelerates its fibrillization and enhances the seeding effect. J Biol Chem 301, 108184 (2025). 10.1016/j.jbc.2025.108184

25 Cameron, A. et al. Generation and characterization of a rabbit monoclonal antibody site-specific for tau O-GlcNAcylated at serine 400. FEBS Lett **5**87, 3722–3728 (2013). 10.1016/j.febslet.2013.09.042

26 Sharma, A., Singh, A., Debnath, R., Gupta, G. D. & Sharma, K. Role of O-GlcNAcylation in Alzheimer’s disease: Insights and perspectives. European Journal of Medicinal Chemistry Reports 12, 100195 (2024). 10.1016/j.ejmcr.2024.100195

27 Alhadidy, M. M., Stemmer, P. M. & Kanaan, N. M. O-GlcNAc modification differentially regulates microtubule binding and pathological conformations of tau isoforms in vitro. J Biol Chem 301, 108263 (2025). 10.1016/j.jbc.2025.108263

28 Krebs, M. R., Bromley, E. H. & Donald, A. M. The binding of thioflavin-T to amyloid fibrils: localisation and implications. J Struct Biol 149, 30–37 (2005). 10.1016/j.jsb.2004.08.002

29 Mukrasch, M. D. et al. Structural polymorphism of 441-residue tau at single residue resolution. PLoS Biol 7, e34 (2009). 10.1371/journal.pbio.1000034

30 Nizynski, B., Dzwolak, W. & Nieznanski, K. Amyloidogenesis of Tau protein. Protein Sci 26, 2126–2150 (2017). 10.1002/pro.3275

31 Wegmann, S. et al. Tau protein liquid-liquid phase separation can initiate tau aggregation. Embo j 37 (2018). 10.15252/embj.201798049

32 Majumdar, A., Dogra, P., Maity, S. & Mukhopadhyay, S. Liquid-Liquid Phase Separation Is Driven by Large-Scale Conformational Unwinding and Fluctuations of Intrinsically Disordered Protein Molecules. J Phys Chem Lett 10, 3929–3936 (2019). 10.1021/acs.jpclett.9b01731

33 Boyko, S. & Surewicz, W. K. Tau liquid-liquid phase separation in neurodegenerative diseases. Trends Cell Biol 32, 611–623 (2022). 10.1016/j.tcb.2022.01.011

34 Ambadipudi, S., Biernat, J., Riedel, D., Mandelkow, E. & Zweckstetter, M. Liquid-liquid phase separation of the microtubule-binding repeats of the Alzheimer-related protein Tau. Nat Commun 8, 275 (2017). 10.1038/s41467-017-00480-0

35 Sengupta, P., Balaji, J. & Maiti, S. Measuring diffusion in cell membranes by fluorescence correlation spectroscopy. Methods 27, 374–387 (2002). 10.1016/s1046-2023(02)00096-8

36 Hofmann, H. et al. Polymer scaling laws of unfolded and intrinsically disordered proteins quantified with single-molecule spectroscopy. Proc Natl Acad Sci U S A 109, 16155–16160 (2012). 10.1073/pnas.1207719109

37 Smet-Nocca, C. et al. Identification of O-GlcNAc sites within peptides of the Tau protein and their impact on phosphorylation. Mol Biosyst 7, 1420–1429 (2011). 10.1039/c0mb00337a

38 Gatta, E. et al. Evidence for an imbalance between tau O-GlcNAcylation and phosphorylation in the hippocampus of a mouse model of Alzheimer’s disease. Pharmacol Res 105, 186–197 (2016). 10.1016/j.phrs.2016.01.006

39 Cantrelle, F. X. et al. Phosphorylation and O-GlcNAcylation of the PHF-1 Epitope of Tau Protein Induce Local Conformational Changes of the C-Terminus and Modulate Tau Self-Assembly Into Fibrillar Aggregates. Front Mol Neurosci 14, 661368 (2021). 10.3389/fnmol.2021.661368

40 Avila, J. et al. Tau Phosphorylation by GSK3 in Different Conditions. Int J Alzheimers Dis 2012, 578373 (2012). 10.1155/2012/578373

41 Jayapalan, S. & Natarajan, J. The role of CDK5 and GSK3B kinases in hyperphosphorylation of microtubule associated protein tau (MAPT) in Alzheimer’s disease. Bioinformation 9, 1023–1030 (2013). 10.6026/97320630091023

42 Mondragón-Rodríguez, S., Perry, G., Luna-Muñoz, J., Acevedo-Aquino, M. C. & Williams, S. Phosphorylation of tau protein at sites Ser(396-404) is one of the earliest events in Alzheimer’s disease and Down syndrome. Neuropathol Appl Neurobiol 40, 121–135 (2014). 10.1111/nan.12084

43 Chakraborty, P. et al. GSK3β phosphorylation catalyzes the aggregation of tau into Alzheimer’s disease-like filaments. Proc Natl Acad Sci U S A 121, e2414176121 (2024). 10.1073/pnas.2414176121

44 Gibbons, G. S., Lee, V. M. Y. & Trojanowski, J. Q. Mechanisms of Cell-to-Cell Transmission of Pathological Tau: A Review. JAMA Neurol 76, 101–108 (2019). 10.1001/jamaneurol.2018.2505

45 Zhu, Y. et al. Pharmacological Inhibition of O-GlcNAcase Enhances Autophagy in Brain through an mTOR-Independent Pathway. ACS Chem Neurosci 9, 1366–1379 (2018). 10.1021/acschemneuro.8b00015

46 Yuzwa, S. A. et al. Pharmacological inhibition of O-GlcNAcase (OGA) prevents cognitive decline and amyloid plaque formation in bigenic tau/APP mutant mice. Mol Neurodegener 9, 42 (2014). 10.1186/1750-1326-9-42

47 Riba, M. et al. Uncovering tau in wasteosomes (corpora amylacea) of Alzheimer’s disease patients. Front Aging Neurosci 15, 1110425 (2023). 10.3389/fnagi.2023.1110425

48 Jeganathan, S., von Bergen, M., Brutlach, H., Steinhoff, H. J. & Mandelkow, E. Global hairpin folding of tau in solution. Biochemistry 45, 2283–2293 (2006). 10.1021/bi0521543

49 Yuzwa, S. A. et al. Increasing O-GlcNAc slows neurodegeneration and stabilizes tau against aggregation. Nat Chem Biol 8, 393–399 (2012). 10.1038/nchembio.797

50 Bijttebier, S. et al. IP-LC-MSMS Enables Identification of Three Tau O-GlcNAcylation Sites as O-GlcNAcase Inhibition Pharmacodynamic Readout in Transgenic Mice Overexpressing Human Tau. J Proteome Res 22, 1309–1321 (2023). 10.1021/acs.jproteome.2c00822

51 Augustinack, J. C., Schneider, A., Mandelkow, E. M. & Hyman, B. T. Specific tau phosphorylation sites correlate with severity of neuronal cytopathology in Alzheimer’s disease. Acta Neuropathol 103, 26–35 (2002). 10.1007/s004010100423

52 Guillozet-Bongaarts, A. L. et al. Pseudophosphorylation of tau at serine 422 inhibits caspase cleavage: in vitro evidence and implications for tangle formation in vivo. J Neurochem 97, 1005–1014 (2006). 10.1111/j.1471-4159.2006.03784.x

53 Mondragón-Rodríguez, S. et al. Cleavage and conformational changes of tau protein follow phosphorylation during Alzheimer’s disease. Int J Exp Pathol 89, 81–90 (2008). 10.1111/j.1365-2613.2007.00568.x

54 Kátai, E. et al. Oxidative stress induces transient O-GlcNAc elevation and tau dephosphorylation in SH-SY5Y cells. J Cell Mol Med 20, 2269–2277 (2016). 10.1111/jcmm.12910

55 Liu, M. et al. Hyperphosphorylation Renders Tau Prone to Aggregate and to Cause Cell Death. Mol Neurobiol 57, 4704–4719 (2020). 10.1007/s12035-020-02034-w

56 Ercan, E. et al. A validated antibody panel for the characterization of tau post-translational modifications. Mol Neurodegener 12, 87 (2017). 10.1186/s13024-017-0229-1

57 Hanger, D. P. et al. Novel phosphorylation sites in tau from Alzheimer brain support a role for casein kinase 1 in disease pathogenesis. J Biol Chem 282, 23645–23654 (2007). 10.1074/jbc.M703269200

58 Fowler, S. L. et al. Tau filaments are tethered within brain extracellular vesicles in Alzheimer’s disease. Nat Neurosci 28, 40–48 (2025). 10.1038/s41593-024-01801-5

59 Tiernan, C. T. et al. Pseudophosphorylation of tau at S422 enhances SDS-stable dimer formation and impairs both anterograde and retrograde fast axonal transport. Exp Neurol 283, 318–329 (2016). 10.1016/j.expneurol.2016.06.030

60 Saman, S. et al. Exosome-associated tau is secreted in tauopathy models and is selectively phosphorylated in cerebrospinal fluid in early Alzheimer disease. J Biol Chem 287, 3842–3849 (2012). 10.1074/jbc.M111.277061

61 Merezhko, M. et al. Secretion of Tau via an Unconventional Non-vesicular Mechanism. Cell reports 25, 2027–2035.e2024 (2018). 10.1016/j.celrep.2018.10.078

62 Simón, D., García-García, E., Royo, F., Falcón-Pérez, J. M. & Avila, J. Proteostasis of tau. Tau overexpression results in its secretion via membrane vesicles. FEBS Lett 586, 47–54 (2012). 10.1016/j.febslet.2011.11.022

63 Pooler, A. M., Phillips, E. C., Lau, D. H., Noble, W. & Hanger, D. P. Physiological release of endogenous tau is stimulated by neuronal activity. EMBO Rep 14, 389–394 (2013). 10.1038/embor.2013.15

64 Pernègre, C., Duquette, A. & Leclerc, N. Tau Secretion: Good and Bad for Neurons. Front Neurosci 13, 649 (2019). 10.3389/fnins.2019.00649

65 Goedert, M. Cryo-EM structures of τ filaments from human brain. Essays Biochem 65, 949–959 (2021). 10.1042/ebc20210025

66 Passarella, D. & Goedert, M. Beta-sheet assembly of Tau and neurodegeneration in Drosophila melanogaster. Neurobiol Aging 72, 98–105 (2018). 10.1016/j.neurobiolaging.2018.07.022

67 Fitzpatrick, A. W. P. et al. Cryo-EM structures of tau filaments from Alzheimer’s disease. Nature 547, 185–190 (2017). 10.1038/nature23002

68 Buschmann, V., Krämer, B., Koberling, F., Macdonald, R. & Rüttinger, S. Quantitative FCS: determination of the confocal volume by FCS and bead scanning with the microtime 200. Application Note PicoQuant GmbH, Berlin (2009).

