## Supplemental files for "*O*-GlcNAcylation effect on Tau in modulating its seeding and cellular transmission in Alzheimer’s Disease"

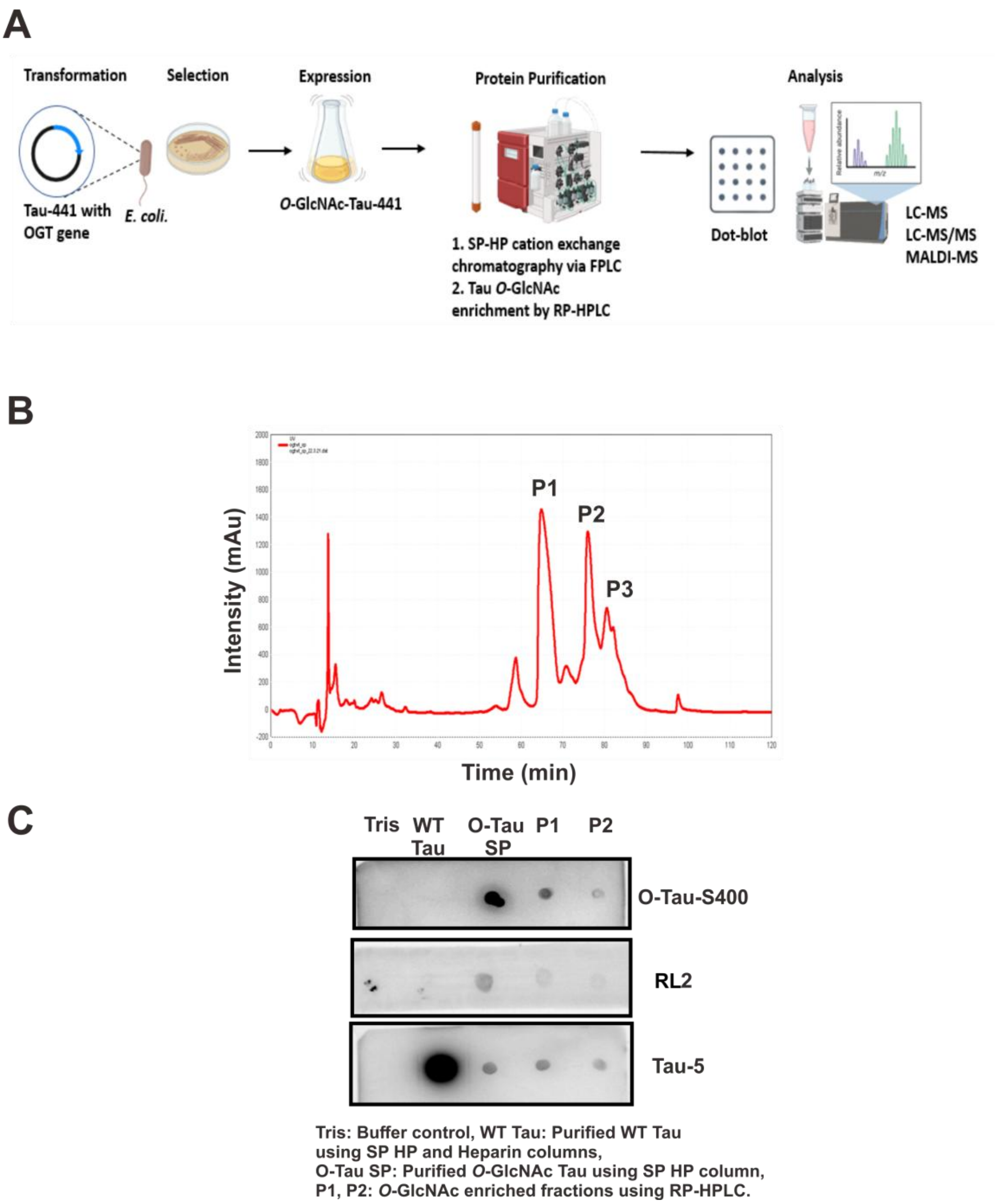


**Figure S1. Purification and characterization of *O*-GlcNAcylated full-length Tau-441. (A)** Schematic representation of the purification and characterization of *O*-GlcNAc-Tau. The recombinant O-Tau was expressed from co-overexpression of Tau-441 with OGT in E. *coli.* culture and purified from SP-HP cation exchange chromatography and RP-HPLC. **(B)** RP-HPLC chromatogram of O-Tau purification. The elution peaks of Tau were labeled as P1, P2, and P3 fraction. **(C)** Dot-blot analysis of the designated P1 and P2 elution fractions obtained from RP-HPLC. The fractions were detected with Tau *O*-GlcNAcylation at S400 site by antibody (anti-*O*-GlcNAc-Tau-S400), a pan-specific antibody for *O*-GlcNAcylation (RL2), and total Tau antibody (Tau-5), confirming Tau *O*-GlcNAcylation.

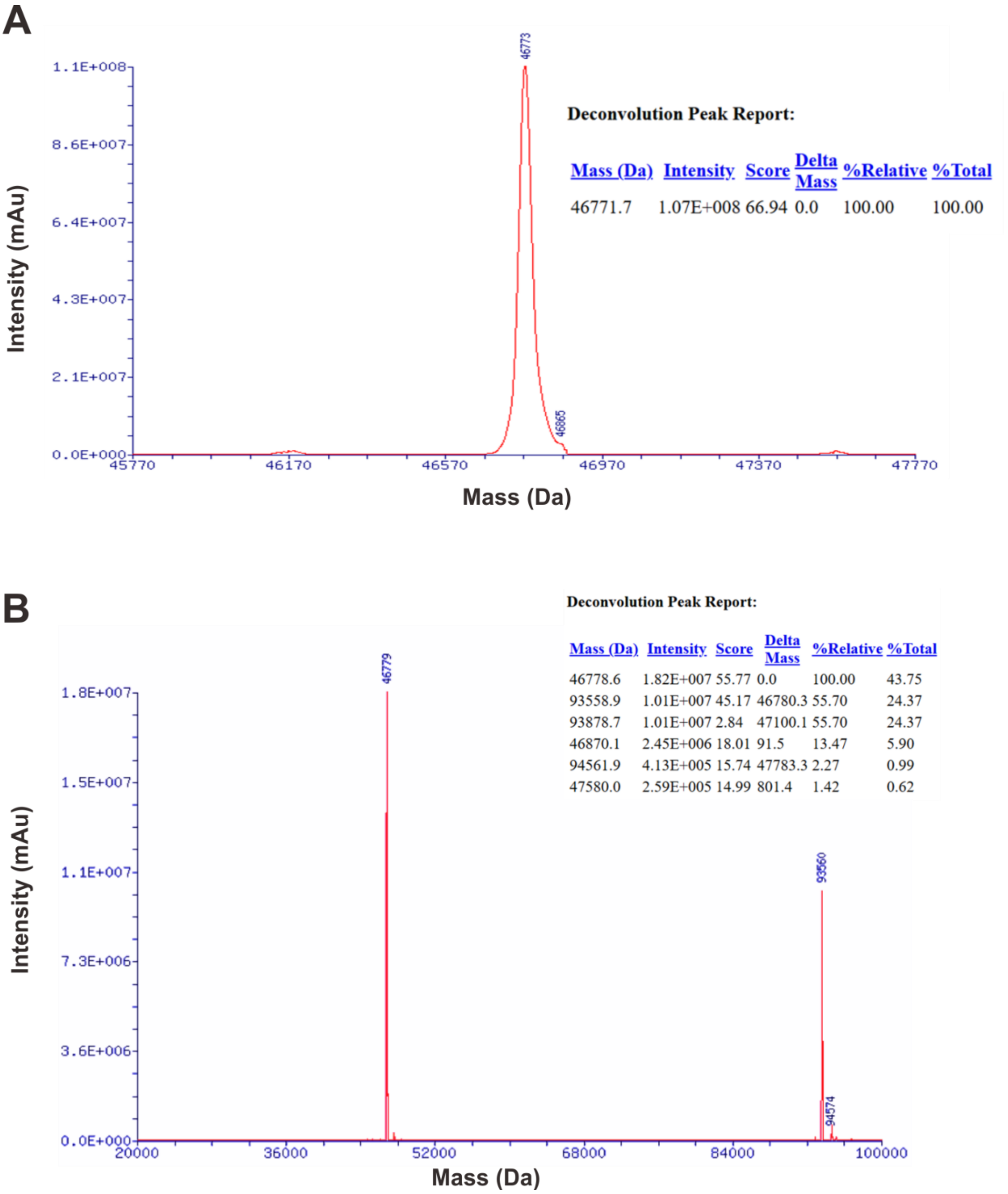


**Figure S2**. **Characterization and determination of *O*-GlcNAcylation on full-length Tau-441 by LC-MS.** LC-MS analysis with deconvolution peak report for P1 fraction **(A)** and P2 fraction **(B)**, respectively. P1 fraction contained monomeric species of O-Tau with an average of ~4-5 O-GlcNAc modifications per Tau molecule. P2 peak was a combination of monomer and dimer species with peaks at ~46.7 kDa and ~93.5 kDa, respectively.

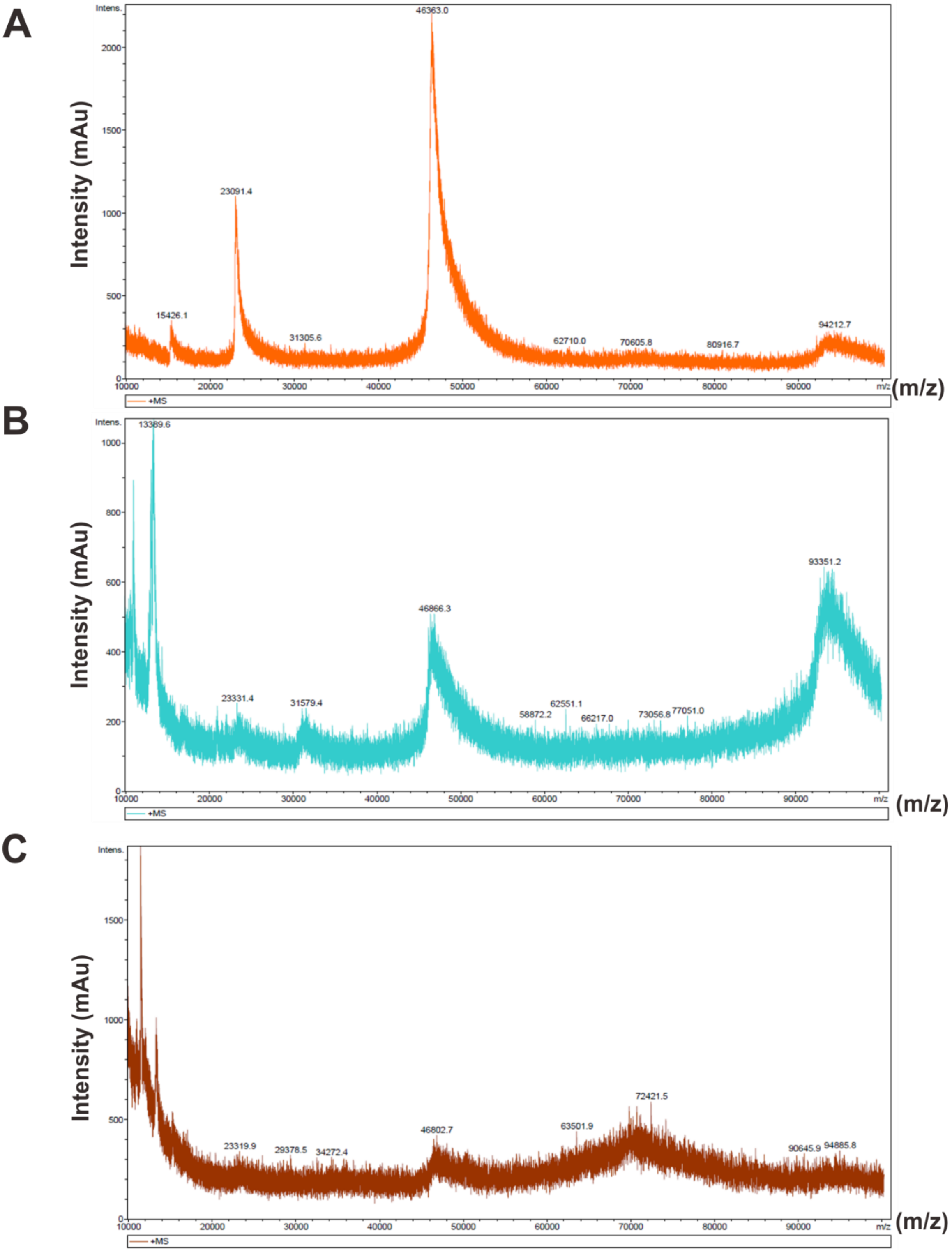


**Figure S3. Characterization of *O*-GlcNAcylation on full-length Tau-441 by MALDI-MS** (**A-C**) MALDI-TOF mass spectra for the elution fractions P1 **(A)**, P2 **(B)**, P3 **(C)** respectively, obtained after *O*-GlcNAc enrichment by RP-HPLC. P1 mass data showed a high intensity predominantly monomeric peak at m/z 46.3 kDa, conforming an average of ~4 *O*-GlcNAc modifications per Tau molecule, and P2 having a dimeric peak of 93.3 kDa besides the monomer peak, and P3 being mostly noise.

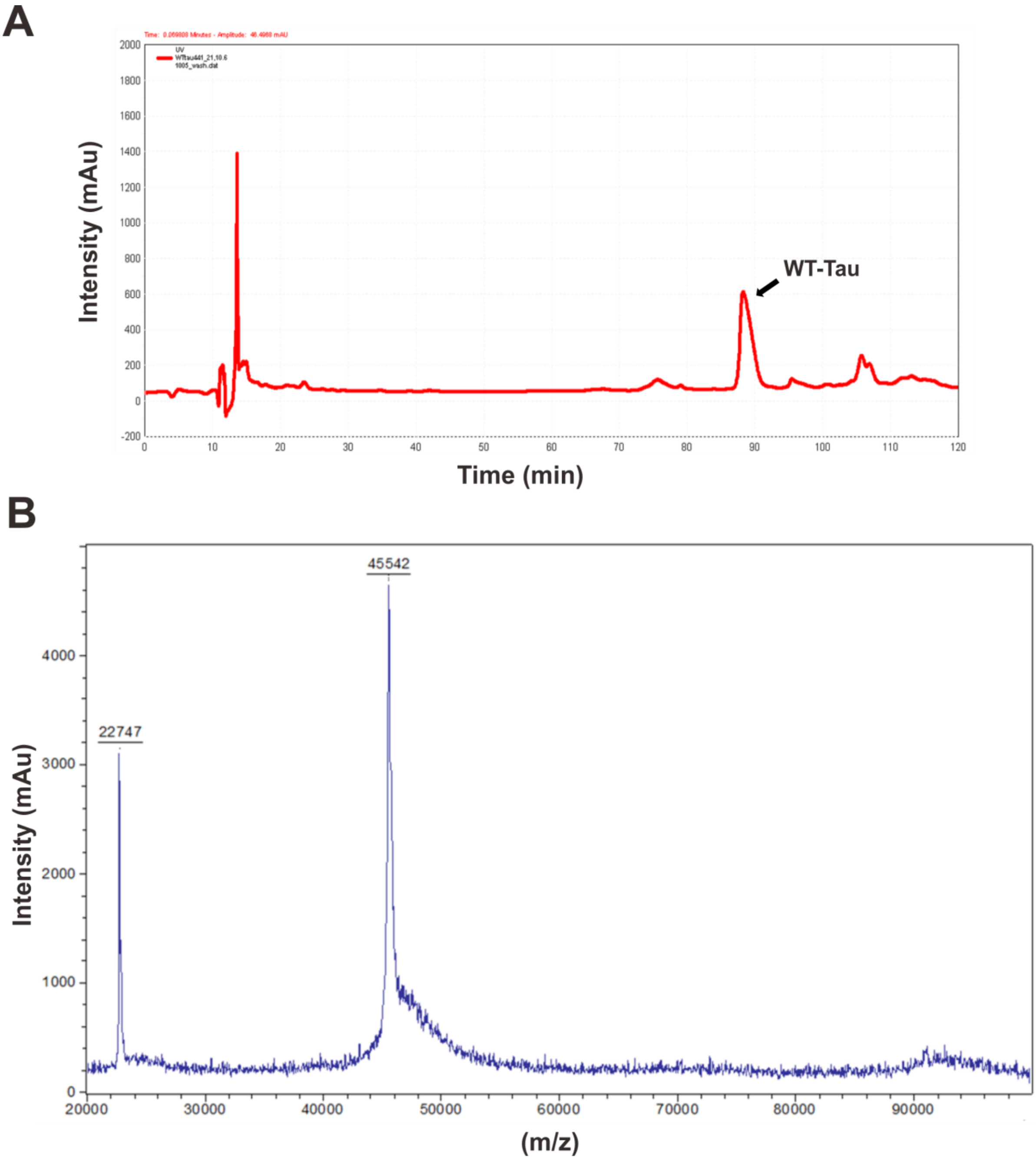


**Figure S4. Characterization of WT Tau. (A)** Chromatogram for WT Tau using the same purification procedure for O-Tau. A single elution peak around 90 mins was detected. **(B)** MALDI-TOF mass analysis detected a monomeric peak at 45.5 kDa for unmodified WT-Tau-441.

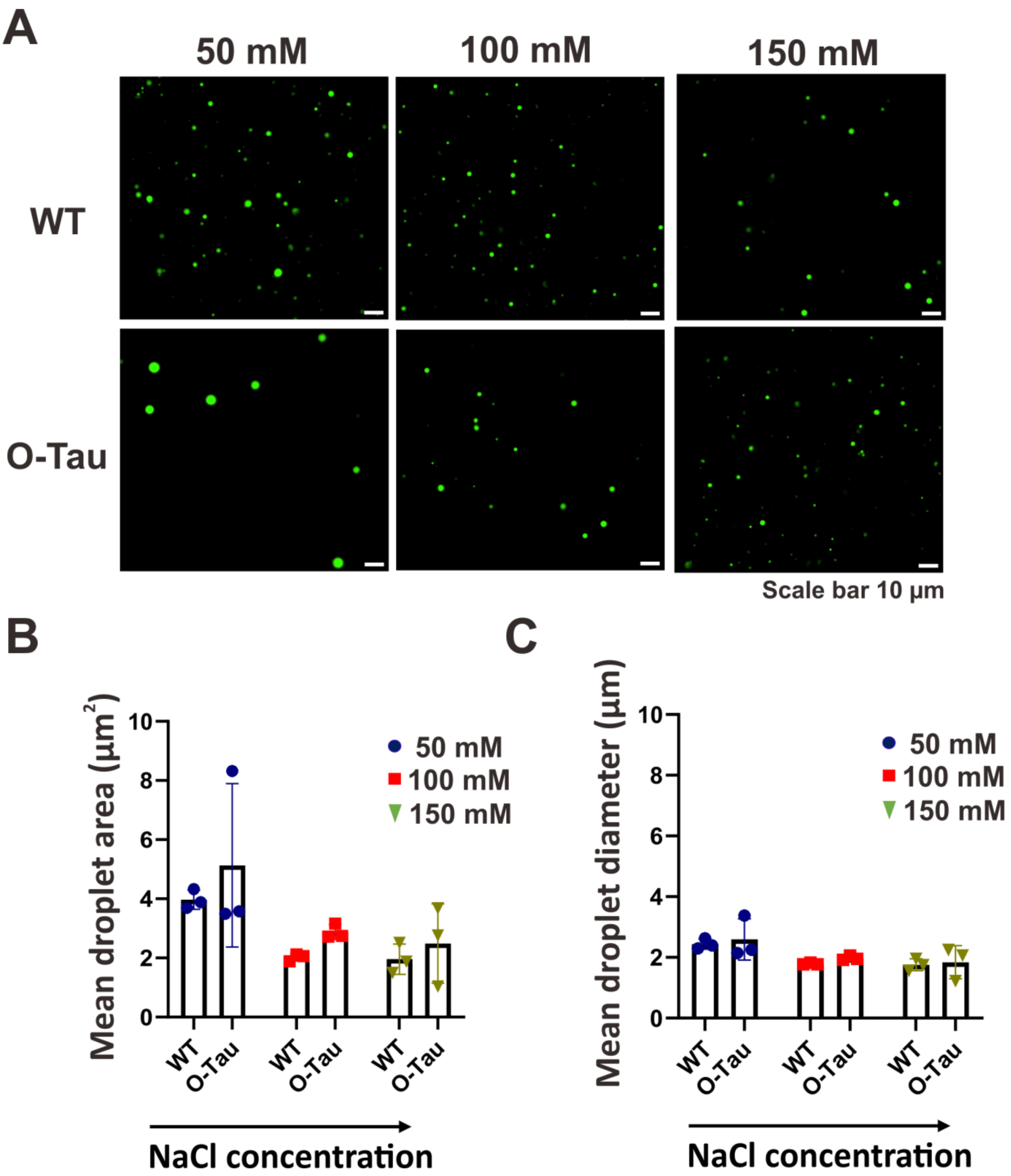


**Figure S5. Droplet induction of Tau by LLPS**. **(A)** Confocal fluorescence microscopy imaging of Alexa flour 488 labelled 10 µM WT and O-Tau monomer. Spherical droplets were detected with 10 % PEG 8000 in HEPES buffer pH 7.4 at 50 mM, 100 mM and 150 mM of NaCl s respectively, (Scale bar= 10 µm). **(B-C)** Quantification of the droplets. The plots for mean droplet area **(B)** and diameter **(C)** of the droplets with increasing concentration of NaCl are shown (n=3).

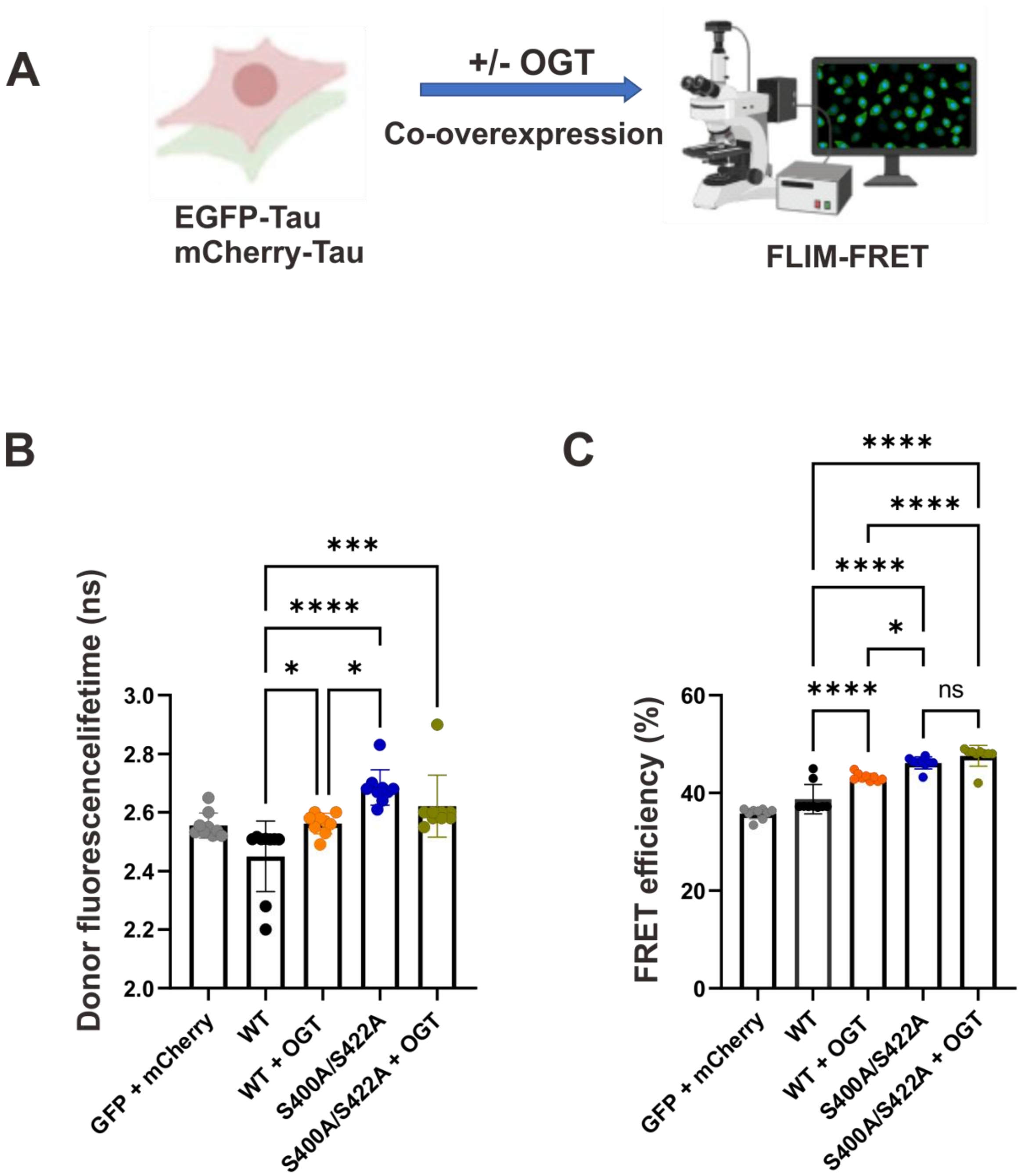


**Figure S6: FLIM-FRET analysis of Tau in live HEK293 cells. (A)** Schematic representation of co-transfection of EGFP-WT and m-Cherry-WT and S400A/S422A double mutant with or without OGT overexpression followed by FILM-FRET analysis after 48 hr incubation. Quantification of donor fluorescence lifetimes are shown in **(B)** and % FRET efficiencies are shown in **(C).** Data are presented as mean ± (s.d.); n=9 cells per condition. Statistical significance was calculated by one-way ANOVA, with Tukey's multiple comparison post hoc test. **P<0.05, **P<0.01, ***P<0.001*.

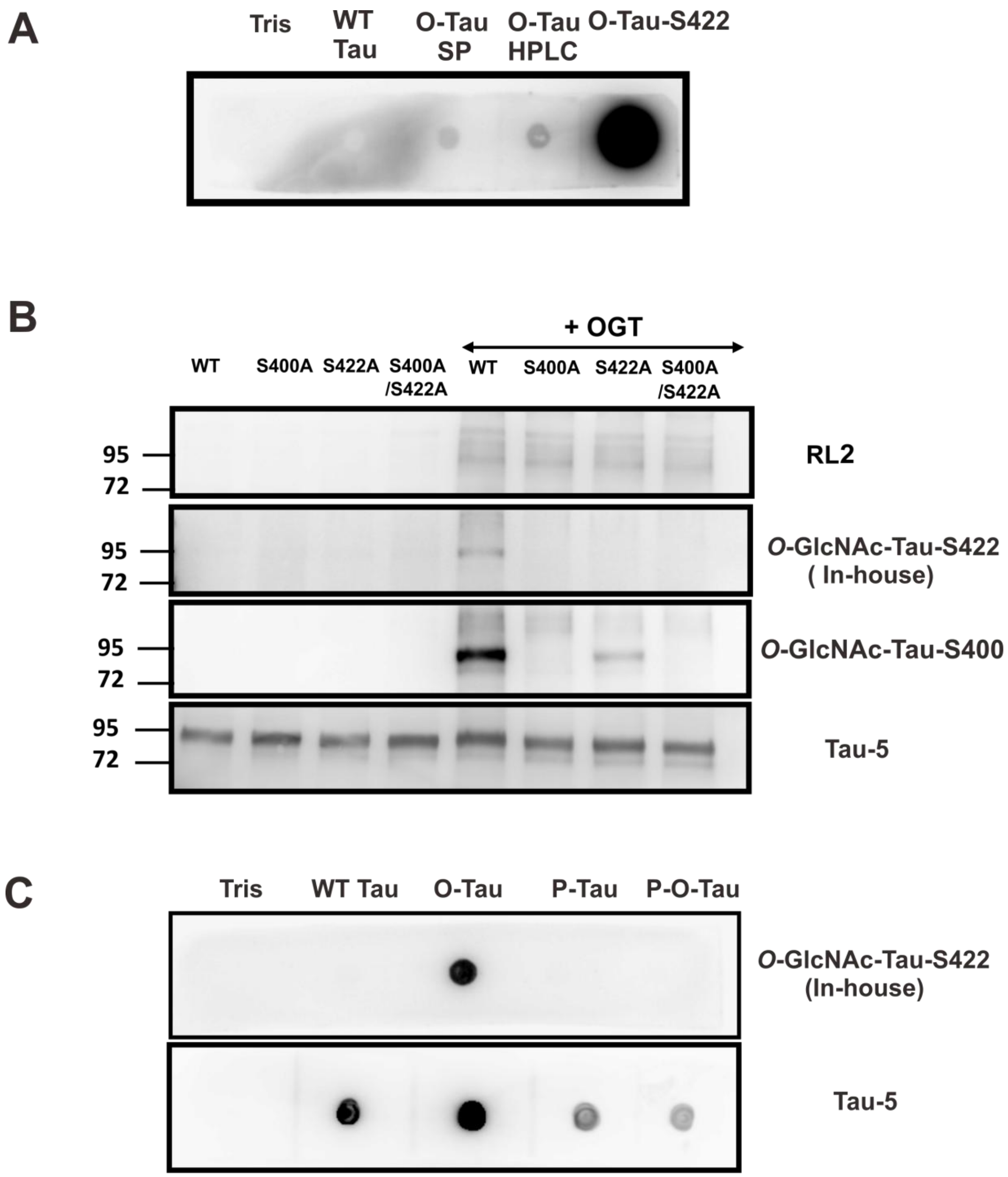


**Figure S7. Characterization of in-house *O*-GlcNAc-Tau-S422 antibody in vitro**. **(A)** Dot-blot analysis of S422 *O*-GlcNAcylation in recombinantly expressed Tau. Several Tau samples were dotted including Tris buffer. The recombinantly purified *O*-GlcNAc-Tau by SP-HP cation exchange column in Tris buffer (O-Tau SP), the *O*-GlcNAc Tau enrichment by RP-HPLC after SP-HP purification in HEPES buffer (O-Tau-HPLC), and Tau-(412-430)-O-GlcNAc-S422 peptide, which is the peptide containing amino acid residues (412-430) with S422 site specific *O*-GlcNAc, in HEPES buffer (Tau-O-S422). The peptide serves as a positive control. **(B)** Western blot analysis of EGFP WT (gT-WT), S400A, S422A (single) and S400A/S422A (double mutants), respectively, with or without

OGT overexpression. Tau was immunoprecipitated by total Tau antibody Tau-5 and blotted against *O*-GlcNAc-Tau-S422 (in-house) antibody, *O*-GlcNAc-Tau-S400 (Anaspec, commercial antibody), RL2 antibody and Tau-5 antibodies, respectively. **(C)** Dot-bot analysis of Tris buffer control, recombinant WT Tau, O-Tau monomers with GSK-3ß kinase (P-Tau) and without GSK-3ß kinase (P-O-Tau) incubation with *O*-GlcNAc-Tau-S422 (in-house) and Tau-5 antibodies, respectively.
